# Comparative population genomics of relictual Caribbean island *Gossypium hirsutum*

**DOI:** 10.1101/2025.06.02.657498

**Authors:** Weixuan Ning, Guanjing Hu, Daojun Yuan, Mark A Arick, Chuan-Yu Hsu, Zenaida V Magbanua, Olga Pechanova, Daniel G Peterson, Yating Dong, Joshua A Udall, Corrinne E Grover, Jonathan F Wendel

## Abstract

*Gossypium hirsutum* is the world’s most important source of cotton fiber, yet the diversity and population structure of its wild forms remain largely unexplored. The complex domestication history of *G. hirsutum* combined with reciprocal introgression with a second domesticated species, *G. barbadense,* has generated a wealth of morphological forms and feral derivatives of both species and their interspecies recombinants, which collectively are scattered across a large geographic range in arid regions of the Caribbean basin. Here we assessed genetic diversity within and among populations from two Caribbean islands, Puerto Rico (n = 43, five sites) and Guadeloupe (n = 25, one site), which contain putative wild and/or introgressed forms. Using whole-genome resequencing data and a phylogenomic framework derived from a broader genomic survey, we parsed individuals into feral derivatives and truly wild forms. Feral cottons display uneven levels of genetic and morphological resemblance to domesticated cottons, with diverse patterns of genetic variation and heterozygosity. These patterns are inferred to reflect a complex history of interspecific and intraspecific gene flow that is spatially highly variable in its effects. Wild cottons in both Caribbean islands appear to be relatively inbred, especially the Guadeloupe samples. Our results highlight the dynamics of population demographics in relictual wild cottons that experienced profound genetic bottlenecks associated with repeated habitat destruction superimposed on a natural ecogeographical distribution comprising widely scattered populations. These results have implications for conservation and utilization of wild diversity in *G. hirsutum*.

## INTRODUCTION

Most modern crop plants have undergone thousands of years of domestication, during which human-mediated directional selection has led to variously severe genetic bottlenecks associated with the favoring of desired morphological traits (Purugganan and Fuller 2009; Andersson and Purugganan 2022; Alam and Purugganan 2024). In contrast, hybridization and introgression, both as natural processes and during crop breeding, serve to increase genetic diversity through combining genetic pools from related crop species (e.g., Katche et al. 2019). These two countervailing processes complicate our ability to unravel the history and genomic structure of modern cultivated species (Gepts 2014). Insights into the domestication history and molecular changes in crop plants thus require analyses of naturally occurring genomic variation across the spectrum, from truly wild species through the various stages spanning original crop domestication to the development of modern cultivars (Olsen and Wendel 2013; Alseekh et al. 2021).

As humans colonized and spread across the Americas, early domesticators became attracted to a number of useful wild plant species (Harlan 1971), including two cotton species (Viot and ^W^endel 2023^)^, *Gossypium hirsutum* (genome designation AD_1_) and *G. barbadense* (AD_2_). These species have seeds with special single-celled epidermal seed trichomes, colloquially known as cotton “fibers”, which are short and light brown in their natural wild forms. Human utilization and domestication of these two wild species has a long history, over 7,000 years in *G. barbadense* and over 4,000 years in *G. hirsutum* (Wendel et al. 1992; Brubaker and Wendel 1994; Brubaker et al. 1999; Westengen et al. 2005; Splitstoser et al. 2016). This long period of protracted domestication and subsequent improvement occurred in parallel in these two species, which in the wild exist in complete allopatry, with *G. barbadense* originating in South America and *G. hirsutum* in Central America (reviewed in Viot and Wendel 2023). In both cases, strong human-mediated selection transformed the wild plant architecture from perennial, multi-branched outcrossing shrubs bearing sparse, relatively short fibers into modern, high-yielding annualized selfing crop plants bearing the strong, fine, long fibers that form the basis of the contemporary global cotton trade (Wendel et al. 1992).

Modern cultivars of *G. hirsutum* presently account for the majority of cotton produced globally. Despite this agricultural significance, little is known about the genetic diversity present within the wild forms. This knowledge gap limits our ability to fully elucidate the consequences and mechanisms of domestication and constrains the effective use of genomic resources for crop improvement (Hu et al. 2021). As with many other important crop plants, truly wild forms of *G. hirsutum* are relatively rare (Fryxell 1979; d’Eeckenbrugge and Lacape 2014), and these can be easily confused with both (1) early domesticated (semi-domesticated) forms that variously overlap in their ‘wild’ cotton characteristics (Fryxell 1979; Brubaker and Wendel 1994; Brubaker et al. 1999; Yuan et al. 2021), and (2) feral derivatives that escaped from cultivation over the millennia and became reestablished in various parts of the presumed ancestral species range (Fryxell 1979; Wendel et al. 1992; Brubaker and Wendel 1994; Brubaker et al. 1999; Alavez et al. 2021). These semi-domesticated and feral forms of *G. hirsutum* are quite common throughout the historical and modern cotton growing regions of the subtropics, including much of Central America, the Caribbean, and northern South America. Because of the complexities introduced by protracted domestication across the millenia, geographic diffusion throughout much of the tropical and subtropical Americas, and repeated escape from domestication as reestablished feral populations, the nature of what constitutes truly wild cotton has been fraught and widely discussed (Hutchinson 1951; Fryxell 1979; Wendel et al. 1992; Brubaker and Wendel 1994; Brubaker et al. 1999; d’Eeckenbrugge and Lacape 2014). This complexity notwithstanding, some efforts have been made to organize this diversity into geo-morphological clusters, for example, the seven “races” of wild or feral cotton described by Hutchinson (1951).

Compounding the foregoing confusion as to what comprises truly wild populations, the parallel domestication of *G. barbadense* and *G. hirsutum* was accompanied by similar processes of geographic diffusion and escape into nature as reestablished feral derivatives (Percy and Wendel 1990; Wendel et al. 1992; Wang et al. 2019). Although these two species were initially domesticated in different continents, i.e., in the Yucatan Peninsula for *G. hirsutum* (Wendel et al. 1992; Brubaker and Wendel 1994; d’Eeckenbrugge and Lacape 2014) and northwest South America for *G. barbadense* (Percy and Wendel 1990; Westengen et al. 2005), they came into contact as a result of thousands of years of human-mediated germplasm diffusion and trade, becoming sympatric in a broad region of the Caribbean and other adjacent locations in northern South America and parts of Central America (Wendel et al. 1992; Brubaker and Wendel 1994; d’Eeckenbrugge and Lacape 2014). This sympatry provided the opportunity for ancient intermingling and introgression (Stephens and Phillips 1972; Brubaker et al. 1993), which now has been documented using extensive whole genome resequencing surveys (Yuan et al. 2021). However, without knowing the genetic composition of truly wild representatives of both species, the inference of introgression is challenging, and thus the extent of introgression between the two species remains unclear.

To shed the light on the nature and extent of the remaining wild cotton genetic diversity, we recently have been collecting *G. hirsutum* from many parts of its indigenous range, with the goal of phylogenomically contextualizing this diversity within a global survey of mostly germplasm bank samples, which we previously completed for the species as a whole (Yuan et al. 2021). Our focus here is on the relictual natural populations of *G. hirsutum* on two Caribbean islands, PuertoRico and Guadeloupe (Fig. 1A). The previous survey (Yuan et al. 2021) suggested that wild cotton may exist near the southwestern coastal tip of Puerto Rico. In addition, Ano el al., (1982) described two types of *G. hirsutum* coexisting in the easternmost part (Pointe des Châteaux) of Guadeloupe (Fig. 1), one with several morphological characteristics that are more typical of *G. barbadense*, and the other, more extensive population with features that characterize wild *G. hirsutum* (Fryxell 1979), specifically the presence of small capsules and seeds, the latter covered with brown to dark green fuzz and khaki-colored fiber.

**Figure 1.**
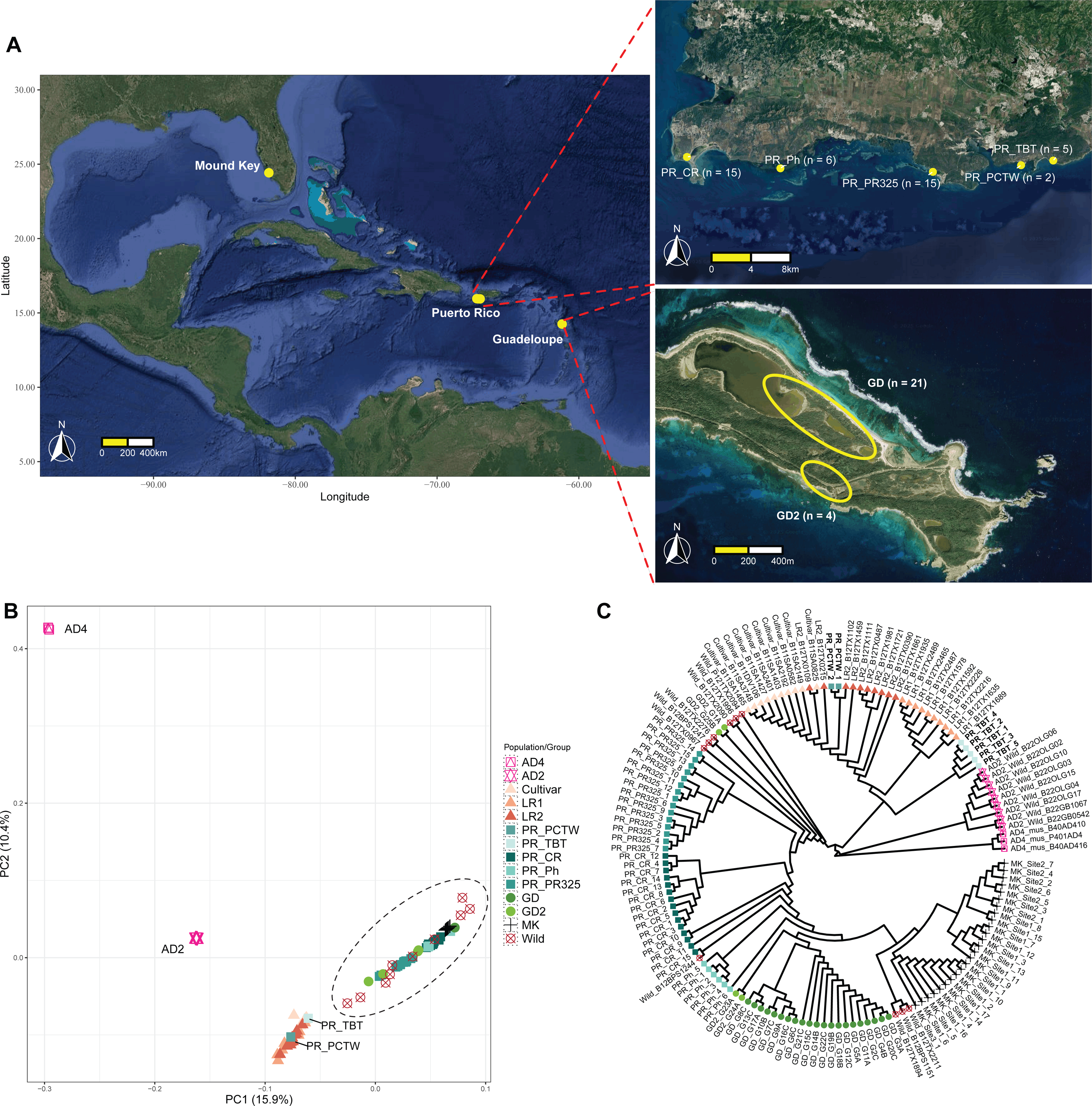
(A) The distribution of field-collected *G. hirsutum* samples from Mound Key, Florida and the Caribbean islands of Puerto Rico and Guadeloupe. Puerto Rican samples (right top) were collected from five sites along the SW coastline, dispersed only a few km between each site (see scale bar). The 25 Guadeloupe samples (right bottom) were genetically partitioned into two groups, GD (n = 21) and GD2 (n = 4), and were collected from one site at Pointe des Châteaux, Guadeloupe. (B) and (C) PCA and neighbor-joining trees showing genetic relationships among Caribbean cottons and *G. mustelinum* (AD4), *G. barbadense* (AD2), Mound Key (MK), and genetic groups from Yuan et al. (2021): Cultivar, Landrace1 (LR1), Landrace2 (LR2), and Wild groups. Maps plotted using R package ggmap (Kahle and Wickham 2013) and resources from Map Data © 2025 Google map (https://www.google.com/maps).

Here we study the newly collected cotton samples from both of these islands using whole genome resequencing of field-collected individuals (n = 25 and n = 43 for Guadeloupe and Puerto Rico, respectively). The sequencing data are analyzed in the context of our previous global survey of germplasm bank accessions (Yuan et al. 2021) and compared to a recently identified wild cotton population from Mound Key, Florida (Ning et al. 2024). We were particularly interested in assessing: (1) whether the *G. hirsutum* populations in Guadeloupe or Puerto Rico are truly wild, or if instead they represent derivatives from some earlier state of cotton cultivation in the Caribbean; (2) how much genetic diversity exists within and between these populations and how different the populations are from each other; and (3) how much heterozygosity is there genome-wide in an average individual, and what this might reveal about population demographic and inbreeding histories.

## MATERIALS AND METHODS

### Whole Genome Resequencing

In March 2018, seeds from 25 *G. hirsutum* individuals were collected by G. A. from the Pointe des Châteaux site of Guadeloupe (Fig. 1A) and sent to Iowa State University where they were planted in the greenhouse (Table S1). Once mature, specimens from each individual were deposited at the Ada Hayden herbarium (ISC; Voucher IDs: from 455774 to 455797). In January 2023, leaves from 43 *G. hirsutum* individuals were collected by J. F. W. from five sites in southeastern Puerto Rico (Fig. 1A), which were immediately dried and stored in silica gel for later DNA extraction.

For the Guadeloupe samples, genomic DNA was extracted from fresh young leaves via Qiagen DNeasy Plant Mini Kit following the manufacturer’s protocol. Extracted DNA concentration and quality were measured using the Qubit™ 1x dsDNA HS Assay Kit on a Qubit™ 4 Fluorometer and NanoDrop™ 2000 Spectrophotometer. DNA libraries were constructed and sequenced by BGI Genomics (Hong Kong) using the BGISEQ-500 system to produce 150 bp paired-end reads.

For the Puerto Rican samples, the silica dried leaf tissues (30 - 50 mg) were ground into fine powder, and genomic DNA was extracted using the Qiagen Plant DNeasy Mini Kit. Purity of the extracted DNA was assessed using a NanoDrop™ One spectrophotometer, and DNA integrity was confirmed via agarose gel electrophoresis. Library preparation used 300 - 500 ng of genomic DNA per sample in the NEBNext Ultra II FS DNA Library Prep Kit with 3-cycle PCR barcoding and enrichment procedure. Genomic libraries were pooled in equimolar amounts and sequenced on a single lane of the Illumina NovaSeq S4 system with 150 bp paired-end reads by Novogene Corporation.

### Experimental Setup and Genome Variant Calling

To determine the wild or domesticated status of cottons from the Caribbean islands, we analyzed 68 individuals (25 Guadeloupe and 43 Puerto Rico) within the context of a global *G. hirsutum* germplasm framework (Yuan et al. 2021). We selected 10 representative individuals each from the inferred natural clusters representing Wild, Landrace1 (LR1), Landrace2 (LR2), and Cultivar groups. In addition, we included all 25 individuals from a recently identified wild *G. hirsutum* population at Mound Key, Florida (Ning et al. 2024). To select the outgroups for genomic introgression tests, we downloaded sequencing data for wild *G. barbadense* (genome designation AD_2_; n = 18) and *G. darwinii* (AD_5_; n = 5) from Yuan et al (2021), and a more distantly related species *G. mustelinum* (AD_4_; n = 3) (Table S1; S2).

Raw reads were trimmed using Trimmomatic V.0.39 (Bolger et al. 2014) to remove sequencing adapters and low-quality bases (‘ILLUMINACLIP:Adapters.fa:2:30:10:2:True LEADING:3 TRAILING:3 MINLEN:75’). Trimmed paired-end reads were mapped to the *de novo* assembled reference genome of wild *G. hirsutum* accession TX2094 v2.0 (unpub. data) via BWA v0.7.17 with flag ‘-R’ to add read group information (Li and Durbin 2009). For samples split over several lanes, multiple bam files were generated, each containing relevant read group information (https://github.com/Wendellab/CaribbeanAD1). The Sentieon DNASeq variant-calling pipeline (Kendig et al. 2019) was applied to genotype variants from each bam file. Specifically, duplicated reads were removed (‘--rmdup’) and small indels were realigned (‘--algo realigner’) to increase precision. From the final realigned bam file, we performed base quality score recalibration (‘—algo QualCal’) and calculated coverage depth (‘--algo CoverageMetrics’), before calling genomic variants (‘--algo Haplotyper --emit_mode gvcf’) for all 159 samples.

Using these gVCF files, we joint-genotyped samples independently using Sentieon (‘--algo GVCFtyper’) in seven groups: (1) Mound Key, (2) Puerto Rico, (3) Guadeloupe, (4) the group of 40 reference *G. hirsutum* germplasm samples described above, and (5,6,7) the three outgroup species. For each VCF, we filtered for biallelic SNPs with average read coverage between 10 to 100 reads per site (‘--remove-indels --max-missing-count 0 --max-alleles 2 --min-meanDP 10 -- max-meanDP 100 --mac 2’) and invariant sites with the same read coverage (‘--remove-indels -- max-maf 0 --min-meanDP 10 --max-meanDP 100’) via VCFtools v0.1.16; filtered variant and invariant sites were then remerged (‘concat’) via bcftools v1.19 (Li 2011) to create the final VCF for each group.

### Identification of Outgroups and Feral Cottons

To select *G. barbadense* accessions most likely to be truly wild among the 18 sequenced samples previously identified as putatively wild (Yuan et al. 2021), and acknowledging the possible genomic introgression from *G. hirsutum* into both wild *G. barbadense* and *G. darwinii* (Wendel and Percy 1990; Yuan et al. 2021), we first combined the VCFs from Mound Key, *G. darwinii*, *G. barbadense* and *G. mustelinum* (‘merge’) and refiltered the biallelic sites (‘-m2 -M2 -i ’F_MISSING=0’ -q 0.001:minor’) via bcftools (8,090,831 sites). SNP sites with high levels of pairwise linkage disequilibrium (LD) were removed (i.e., LD-pruned SNPs) via PLINK v.1.9 (‘-- indep-pairwise 50 10 0.1’) (Purcell et al. 2007). The LD-pruned VCF (520,520 sites) was used to explore population genetic structure via Principal Component Analysis (PCA) (‘--pca 20’) and neighbor-joining tree analysis (‘--distance square 1-ibs’) using PLINK and the ape v.5.8.1 package (Paradis et al. 2004) in R v4.4.1. Using the same set of filtered SNPs, we analyzed population genetic structure using LEA v3.18.0 (Frichot and François 2015), with K (the number of ancestral populations) ranging from 1 to 10, with each K replicated ten times. Because *G. darwinii* is sister to *G. barbadense* (Grover et al. 2015), we required the selected wild *G. barbadense* to be both monophyletic in the neighbor-joining tree and sister to *G. darwinii* (details see Fig S1; Table S2), leaving nine remaining *G. barbadense* samples.

VCFs containing all 145 samples *G. hirsutum* samples (Table S1; S2) from Puerto Rico, Guadeloupe, and Mound Key, as well as the four inferred groups from Yuan et al (2021), were merged with those nine wild *G. barbadense* and three *G. mustelinum* accessions. This combined dataset was used to evaluate the *G. hirsutum* samples for potentially cryptic feral representatives among the Caribbean cottons. Using the same steps as above, we filtered the 35,453,021 biallelic SNPs from the joined VCF to retain only 4,200,388 high-confidence, LD-pruned sites for subsequent PCA and neighbor-joining tree analyses. Results were plotted in R using ggplot2 v3.5.1 (Wilkinson 2011) and ggtree v3.14.0 (Yu et al. 2017).

After identifying two Caribbean cotton populations (PR_PCTW and PR_TBT) as putatively non-wild feral derivatives due to their placement among domesticated *G. hirsutum* groups, we removed those two populations (7 individuals) from the analysis of wild population diversity along with the 40 germplasm collection samples from Yuan et al (2021), which are of less certain provenance. This subset of 86 samples, representing only putatively wild cotton populations, was extracted via bcftools (‘view -S -m2 -M2 -i ’F_MISSING=0’ -q 0.001:minor’) from the combined VCF of 145 samples, to retain only the Mound Key cottons, the Guadeloupe cottons, and the remaining non-feral Puerto Rican cottons. Relationships among these putative populations were evaluated by PCA using the surviving 1,039,022 filtered SNPs, with the first three PCs visualized via a modified script 3D-PCA-plot.py (https://github.com/Siavash-cloud/3D-PCA-plot). In addition, we calculated genetic relatedness among these 86 samples using the PLINK identity-by-descent test (‘--genome’) and plotted the PI_HAT values higher than 0.3) with ggplot2.

### Genomic Introgression Tests

To evaluate potential gene flow among *G. hirsutum* populations and reciprocal genomic introgression with *G. barbadense*, a dataset of 128 individuals represented by 33,774,815 sites containing 3,292,507 independent biallelic SNPs (‘--indep-pairwise 50 10 0.1’) was extracted from the full dataset of the combined 145 samples and 35 million SNPs described above. The selected samples included 25 Mound Key, 25 Guadeloupe, 36 wild Puerto Rico (excluding PR_PCTW and PR_TBT), 30 of the 40 *G. hirsutum* samples (omitting the 10 wild samples) based on Yuan et al. (2021), 9 wild *G. barbadense*, and 3 *G. mustelinum* samples.

Using the ∼3.3 million filtered biallelic SNPs, we first estimated the number of ancestral populations (K= 1 to 18) via LEA, with each K replicated 10 times. The best K was selected based on the lowest cross-entropy value (Frichot and François 2015). From these results and using geographic considerations, we defined ten groups/populations among the sampled *G. hirsutum* individuals.

Gene flow was estimated using the maximum likelihood-based approach implemented in TreeMix v1.13 (Pickrell and Pritchard 2012) by first calculating the allele frequency at each of the 3.3 million SNP sites in PLINK (--freq) for each population/group/species. For TreeMix, the four *G. mustelinum* individuals was assigned as outgroup (‘-root’) and a block size of 1000 SNPs per window (‘-k’) with bootstrap replications (‘-bootstrap’) was used (https://github.com/Wendellab/CaribbeanAD1). In addition, each ‘m’ (i.e., number of estimated gene flow) had five replicate runs, and the best migration model was selected using the R package OptM (Fitak 2021). The final network was visualized using the R script “plotting_funcs.R” in TreeMix.

Putative genomic introgression was evaluated via the ABBA-BABA (or D-statistic-based) method using Dsuite (Malinsky et al. 2021). Specifically, the topology of the maximum likelihood TreeMix non-network was rooted using the outgroup *G. mustelinum* (‘-k 1000’, ‘-root’) and with the final filtered 3.3 million SNPs as input, we calculated the tree branches that showed strong introgression signals (‘Fbranch’) between ten *G. hirsutum* populations/groups and/or *G. barbadense*.

### Phylogenetic Analysis of Plastomes

The conserved and uni-parentally inherited whole chloroplast DNA (i.e., the plastome) has been widely used in cotton comparative genomics (Wendel and Albert 1992; Chen et al. 2017; Wu et al. 2018; Yan et al. 2024). We *de novo* assembled plastomes using GetOrganelle v1.7.7.1 (Jin et al. 2020) (‘get_organelle_from_reads.py’), with default settings, for all 159 samples in this study. These assemblies, along with 339 previously published cotton plastome sequences (Yan et al. 2024) were first aligned via MAFFT v7.508 (Katoh and Standley 2013) to check assembly quality. One sample (AD5_dar_BYU50004) had a large number of mismatch sites, likely due to contamination, and hence was removed from further analysis.

For plastome annotation, one Guadeloupe sample (GD_G7C) was selected and annotated with an online tool GeSeq (Tillich et al. 2017), using a *G. tomentosum* reference sequence (NC_016690) as a basis (Xu et al. 2012). The resulting annotation file was used to transfer annotations to remaining 157 aligned samples via PAG (Qu et al. 2019). Using a customized script (https://github.com/Wendellab/CaribbeanAD1), we then extracted 110 chloroplast gene sequences present across all 158 individuals, including 81 protein coding genes and 29 tRNA coding genes. Each gene was aligned via MAFFT, and all gap sites (‘-nogaps’) were removed via trimAl v1.5 (Capella-Gutiérrez et al. 2009). We then concatenated (‘concat’) all trimmed gene sequences into one sequence via SeqKit v2.9.0 (Shen et al. 2016).

We constructed cpDNA-based phylogenies for these 158 samples using (1) an alignment of the whole plastome, with the internal repeat B (IRB) region removed using the annotation file of GD_G7C and SeqKit, and gap sites trimmed by trimAl; and (2) a concatenated sequence of the 110 genic loci. Maximum Parsimony trees were constructed using PAUP* v4 (Swofford 1998) and employing a Heuristic search with random addition and tree-bisection-reconnection (TBR) branch-swapping. Given the slow evolutionary rate of cpDNA, we also tabulated the number of parsimony informative sites supporting the backbone of the phylogenies in PAUP* using ‘describeTrees’. In addition, we calculated a haplotype network via PopArt v1.7 (Leigh and Bryant 2015) using the TCS method (Templeton et al. 1992) for both datasets. Complementing the above analyses, the same datasets were used for maximum likelihood phylogenetic reconstructions using IQTree2 (Minh et al. 2019) with bootstrapping (‘-B 1000’).

### Estimation of Genomic Diversity

To quantify genomic diversity within and among populations, we first excluded five outlier samples that did not group with other samples collected from the same site (four in GD2 and one PR_Ph_6; see Results) on the PCA. VCFs containing both variant and invariant sites for the remaining 121 samples from the Caribbean, Mound Key, and the four germplasm groups were merged via bcftools. These samples were divided into nine groups based on genetic relationships inferred by LEA (see Results). For each group, genome-wide nucleotide diversity (π) was calculated using pixy v1.2.10 (Korunes and Samuk 2021), with a sliding-window size of 10 kbp. To ensure that comparisons between different populations/groups were not affected by sample size differences, we randomly downsampled each population/group to five individuals (matching the smallest group, Puerto Rico PR_Ph, n = 5) and replicated this downsampling 20 times. Final π values were calculated by counting the average values of each sliding window across all 20 replicates. Using the same downsampling and window setting, pairwise differences of sequence diversity (d_xy_) and population genetic differentiation (F_st_) among the nine populations/groups were also calculated in pixy.

### Assessment of Homozygosity, Heterozygosity, and Linkage Disequilibrium

The length and frequency of long uninterrupted homozygous sites (i.e., identical haplotypes) can be informative with respect to population structure, history, and breeding system (e.g., Ceballos et al. 2018; Kumar et al. 2020). To explore this in cotton, long runs of homozygosity (ROH) were identified for each of 121 samples using 26,763,108 filtered biallelic SNPs (described above) via the ‘slidingRUNS.run’ function in the R package detectRUNS v0.9.6 (Biscarini et al. 2018). This method applied sliding window-based detection, similar to PLINK ROH detection (Meyermans et al. 2020), by specifying the sliding window size as 15 (windowSize = 15), the threshold of overlapping windows of the same state to 0.05, the minimum number of SNPs to 10, the maximum number of heterozygous SNPs and the maximum number of missing SNPs in a sliding window both to 1 (maxOppWindow = 1, maxMissWindow = 1), the maximum distance between consecutive SNPs to 1 Mbp, the minimal length of ROH to 0.25 Mbp, and the lowest SNP density per kbps to 0.001. Using a customized R script (https://github.com/Wendellab/CaribbeanAD1), we tabulated ROH into three length categories: 0.25 to 1 Mbp, 1 to 2 Mbp, and larger than 2 Mbp, and calculated the proportion of total ROH lengths compared to the whole genome using the reference TX2094 V2 for each individual, i.e., the ROH inbreeding coefficients (F_ROH_).

The proportion of heterozygous sites among the 26 million SNPs were estimated using VCFtools (‘--het’) for the 121 samples. Correlations between linkage disequilibrium (*r*^2^) and all pairs of SNP physical distance (in bp) were estimated for the nine groups using the same set of SNPs and PopLDdecay (Zhang et al. 2019). To account for the sample size differences in LD decay comparison, we randomly downsized each population to five individuals (PR_Ph n =5) and repeated this 20 times, then calculated the average *r*^2^ across all replicates and for each distance window.

### Population Demographic Inference

Whole-genome resequencing data offers the possibility of exploring past demographic processes and/or selection using contemporary allele frequencies (Marchi et al. 2021). Toward that end, population demographics of the four cotton populations that had larger sample sizes and were inferred to be wild (Mound Key, Guadeloupe, and wild Puerto Rican cottons from PR325, and CR; see Results) were first analyzed by calculating the observed folded site frequency spectrum (SFS), Watterson’s theta (*θ*_W_), and Tajima’s *D* (Tajima 1989) via ANGSD (Korneliussen et al. ^2^014^)^ using their final realigned bam files as input. Site allele frequencies were tabulated with ‘angsd’ (with settings ‘-doSaf 1 -doMaf 1 -doMajorMinor 1 -doGlf 3 -uniqueOnly -GL 2 - minMapQ 30 -minQ 20 -minInd 25’) and ‘realSFS’ (withs settings ‘-maxIter 100 -P 20 -fold 1’). Tajima’s *D* and Watterson’s theta were estimated via ‘thetaStat’ with window size (‘-win’) 50,000 bp and step size (‘-step’) 10,000 bp. In addition, the expected SFS under neutrality model was estimated using *θ*_W (Hudson 2015)_.

Changes in effective population size (*Ne*) over evolutionary time can be indicative of bottlenecks or population expansion (Mather et al. 2020), and were estimated here via PSMC (Li and Durbin 2011) by selecting one representative individual (PR_CR_1, PR_PR325_15, PR_Ph_3, MK_Site1_1, GD_G17A; see Results) from each of the five wild cotton populations. The five realigned bam files were used as input to reconstruct a consensus fastq file via samtools and bcftools, which was then converted into PSMCFA format via ‘fq2psmcfa -q20’ and split into smaller segment ‘splitfa’ for bootstrapping. We replicated each PSMC analysis 100 times (with settings ‘-N25 -t15 -r5 -b -p "4+25*2+4+6"’), and the final results were plotted using psmc_plot.pl with mutation rate (‘-u’) 4.56e-9 and generation time (‘-g’) to 2.

## RESULTS

### Inferring Truly Wild G. barbadense as Outgroup

Given that reciprocal introgression is known to have occurred between *G. barbadense* and *G. hirsutum* (Stephens and Phillips 1972; Brubaker et al. 1993; 2021), selecting *G. barbadense* accessions that are known to be truly wild is essential for testing potential genomic introgression from *G. barbadense* into wild *G. hirsutum*. Using 0.5 million independent, whole-genome, biallelic SNPs extracted from 25 Mound Key (wild) *G. hirsutum*, 5 *G. darwinii*, 3 *G. mustelinum*, and 18 *G. barbadense* individuals previously identified as putatively wild (Yuan et al. 2021), we identified nine *G. barbadense* samples that represented a monophyletic clade. Although all wild *G. barbadense* and *G. darwinii* formed one cluster in the first two principal components based on the whole-genome data (39.9% variance explained by PC1 and 25.5% by PC2) (Fig S1A), the ancestral population genetic structure analysis suggested that nine *G. barbadense* individuals (dashed box, Fig S1C) were more uniform, particularly for K=4. These samples were also monophyletic in the neighbor-joining tree (Fig S1B), containing one type of chloroplast DNA (see plastome analysis, below), and were collectively sister to the truly wild, non-domesticated endemic species from the Galapagos Islands, *G. darwinii* (Wendel and Percy 1990; Grover et al. 2015). These samples, accordingly, were used to represent purely wild *G. barbadense*.

### Genetic Structure and Relationships among Major G. hirsutum Genetic Groups

Whole-genome resequencing yielded high coverage for both Guadeloupe (≥46×; average 48×, n = 25) and Puerto Rican samples (≥20×; average 25×, n = 43) (Table S1). Combining the genetic data from the total of 145 samples from the Caribbean (n = 68), Mound Key (n = 25), previously identified germplasm groups (40; Yuan et al. 2021, see methods)n = (40; Yuan et al. 2021, see methods), purely wild *G. barbadense* (n = 9), and the best phylogenetic outgroup to the other allopolyploid *Gossypium* species (n = 3; i.e., *G. mustelinum*), we extracted 4.2 million high-quality, genome-wide, independent biallelic SNPs to assess their genetic relationships.

Principal component analysis (PCA) broadly showed four groups in the first two PCs (Fig. 1B), accounting for 15.9% and 10.4% of the genetic variance, respectively. In addition to the two groups comprising *G. mustelinum* and *G. barbadense*, *G. hirsutum* accessions were divided into two groups, one containing the domesticated cottons comprising Landrace1 (LR1), Landrace2 (LR2), and Cultivars, and the other containing all Wild and Mound Key (MK) cottons (denoted by a dashed circle in Fig. 1B). Thus, there is a primary genetic division between purely wild *G. hirsutum* and other germplasm that has been through a long history of domestication and possible introgression with *G. barbadense.* Notably, eight samples collected from two Puerto Rico sites (PR_PCTW and PR_TBT) grouped with the domesticated cottons, whereas all remaining Puerto Rico (PR) and Guadeloupe (GD) samples nested with the other wild cottons from Mound Key and elsewhere. In addition, compared to Mound Key, all Caribbean cottons collected from both islands exhibited larger variation within the Wild cotton cluster.

A neighbor-joining tree constructed from the same set of 4.2 million SNPs supported a similar depiction and resolution of the four groups: *G. mustelinum*, *G. barbadense*, wild cottons, and domesticated cottons (Fig. 1C). Within domesticated cottons, PR_TBT formed a clade that was sister to all Landrace1 samples, and PR_PCTW was nested with the clade that included interdigitating samples representing groups of Landrace2 and Cultivars. As for wild cottons, only Mound Key, Puerto Rico site PR325 and the main Guadeloupe group (GD, n = 21) were monophyletic. By contrast, Puerto Rican cottons collected from sites CR and Ph were mixed and paraphyletic to each other, and four outlier samples (GD2) from Guadeloupe exhibited two divergent lineages, with one (GD2_G24A with GD2_G23A) sister to the main Guadeloupe group (GD, n = 21), and the other one (GD2_G25B with GD2_G1A) nested among all other wild cottons (Fig. 1A, 1C). These results are interpreted as genetic evidence that the PR_TBT and PR_PCTW populations represent feral derivatives of domesticated cottons that escaped to become reestablished as a component of the native vegetation (Fryxell 1979; Wendel et al. 1992; Alavez et al. 2021), and suggest some admixture among the cottons from different locations.

### Detecting Outliers among Wild Caribbean Cottons

To further explore genetic relationships of the outliers among wild Caribbean cottons, we reconstructed the PCA using approximately one million filtered SNPs from 86 samples of wild cotton from Puerto Rico, Guadeloupe, and Mound Key. The first principal component (Fig 2A) explained 23.8% of the variation and separated Mound Key from the remaining Caribbean cottons. In PC2 (12.01%) and PC3 (8.5%), the Caribbean cottons clustered into three loose groups: (1) the Guadeloupe cotton with a main group (GD, n = 21) and four more diverged outliers (GD2, n=4), (2) the Puerto Rican population PR325, and 3) the remaining group of Puerto Rican cottons (CR and Ph). The four Guadeloupe outlier samples (GD2) were grouped in two sets of two individuals each (GD2_G25B with GD2_G1A, and GD2_G24A with GD2_G23A). In addition, the sample PR_Ph_6 collected from the Puerto Rican Ph site (Fig. 1A) was closer in multivariate space to the Guadeloupe main cotton group compared to the other five PR_Ph samples.

**Figure 2.**
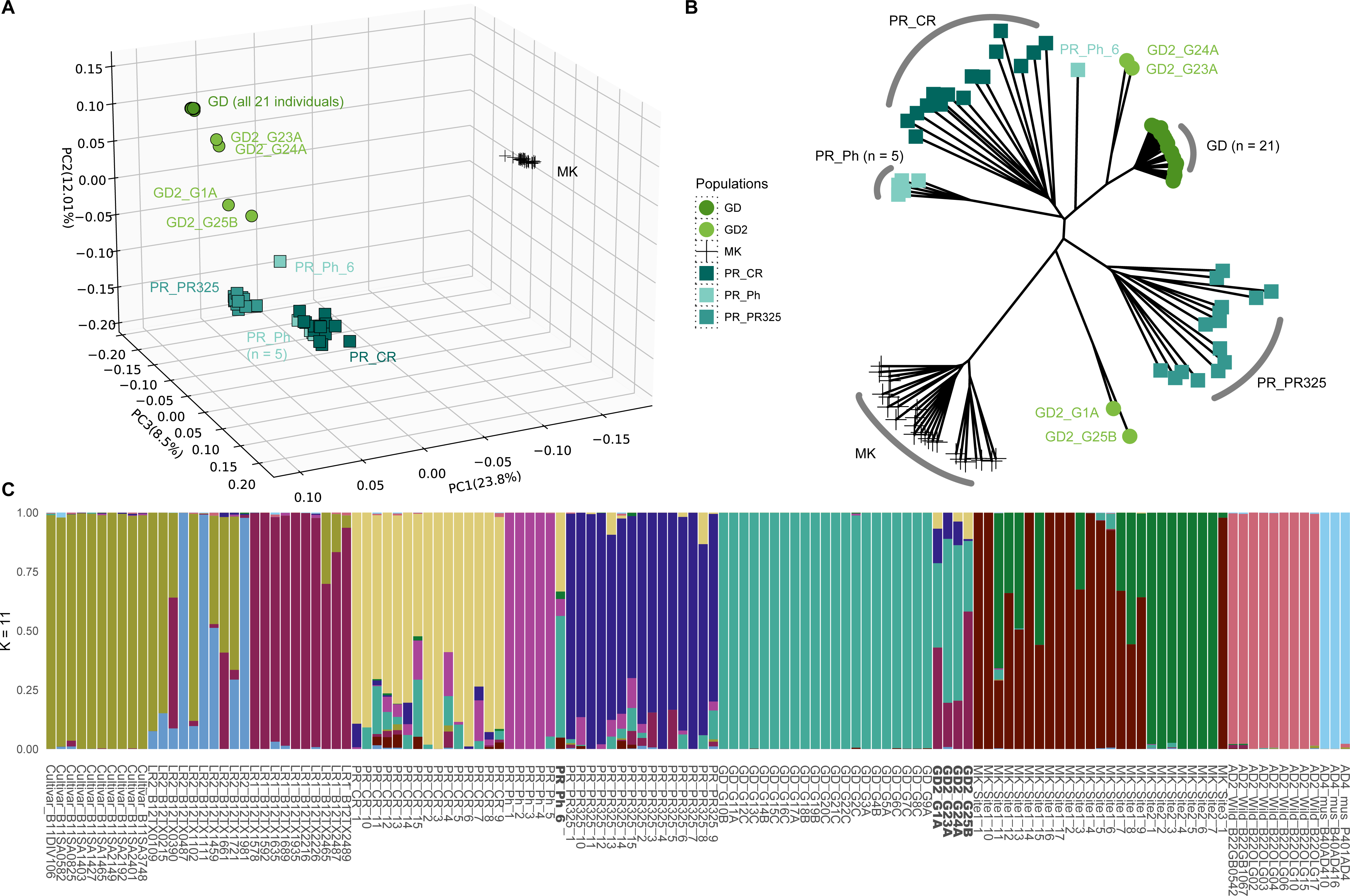
Genetic relationships inferred from (A) PCA and (B) neighbor-joining tree of 61 Caribbean and 25 Mound Key *G. hirsutum* samples. Each accession/population is represented by a different symbol. (C) LEA genetic structure plot when the number of ancestral populations (K) = 11. The x-axis is labeled with individual code (Table S1) and the y-axis represents the proportion of estimated ancestral population and is filled by different colors.

Neighbor-joining analysis (Fig 2B) recapitulates the PCA patterns, whereby most wild cotton populations were monophyletic, with the aforementioned exceptions of PR_Ph_6 and the four Guadeloupe outlier samples (GD2). Specifically, two samples (GD2_G24A and GD2_G23A) formed a separate lineage sister to the Guadeloupe main group (GD, n = 21) and notably adjacent to a lineage composed of solely sample PR_Ph_6, whereas the other two outliers (GD2_G1A and GD2_G25B) formed a lineage that was in between the Mound Key and PR_PR325 clades.

Genetic kinship analysis was explored using the metric PI_HAT for the 86 cotton samples used in this analysis. The relatedness within the main populations of Guadeloupe (GD) and PR_Ph (n = 5) showed mean PI_HAT values of 0.8, respectively, indicating a high level of kinship. This is in contrast to Mound Key cottons whose two geographically close (about 200 m) subpopulations (Ning et al. 2024) displayed an average PI_HAT of 0.57 (Fig. S2). Notably, the two sets of four Guadeloupe outliers exhibited lower levels of relatedness, both with an average PI_HAT of around 0.46. Puerto Rican cottons from the CR and PR325 sites had the lowest relatedness among individuals within sites, being approximately 0.36 and 0.42, respectively.

### Frequent Gene Flow among Caribbean Cottons

To assess the effects of potential inter- and intraspecific gene flow in shaping genetic relationships and diversity of the Caribbean cottons, we included the additional three domesticated *G. hirsutum* gene pools (Landrace1, Landrace2 and Cultivars) and the outgroups (*G. barbadense* and *G. mustelinum*) in a new analysis. In total, 3.3 million independent biallelic SNPs were extracted for a total 128 samples.

Population structure analysis by LEA identified 11 as the optimal number of ancestral populations (K value) (Fig S3), with the results (Fig. 2C) reiterating the inferences from the PCA and NJ phylogenetic analyses (Fig 2A, 2B) described above. Each outgroup species formed its own cluster, as expected, and the domesticated cottons were divided into three clusters, as previously noted (Yuan et al. 2021), with more mixed ancestral signals detected in Landrace2. Among wild cottons, most samples clustered based on their collection sites, except for PR_Ph_6 and four Guadeloupe outliers (GD2). Notably, PR_Ph_6 exhibited over 50% ancestral signals from the Guadeloupe main population (GD), and the four Guadeloupe outliers (GD2) had variable proportions of their ancestry shared with Landrace1, rising to over 50% in GD2_G25B and GD2_G1A. Unlike Puerto Rico Ph (n = 5) and the Guadeloupe (n = 21) main population (GD), which had mostly uniform ancestry, Mound Key cottons exhibited two distinct ancestral populations, as previously reported (Ning et al. 2024). In contrast to these observations of site-specificity, the Puerto Rican cottons from both CR and PR325 exhibited a combination of site-specific major ancestral genetic background with mixed signals from multiple other sources.

Using the same set of SNPs as above, we estimated genomic introgression among all 128 samples using TreeMix (Fig. 3A). We first grouped the samples based on their genetic structure (Fig. 2C), from which we separated the outliers from GD2 and PR_Ph_6 (described above) as two additional groups and merged the two Mound Key subpopulations, for a total of 12 populations/groups. The best TreeMix model estimated the number of migration events (m) = 7 (Fig. S4). Compared to the topology of the maximum likelihood tree when m = 0 (the phylogenetic tree on the left side of the heatmap in Fig. 3B), the topology of the TreeMix result when m = 7 placed four Guadeloupe outliers (GD2) as sister to Landrace1 among all domesticated cotton groups (Fig. 3A), as expected from the above results and indicating mixed ancestry. The best TreeMix model also supported three gene flow branches from Landrace1 into Puerto Rican populations PR325, Ph, and CR, which is also supported by the ancestry structure for PR325 and CR but is not apparent in Ph (Fig. 2C). Two additional gene flow branches with high migration weight were suggested from the Guadeloupe main population (GD) into both the GD2 and Mound Key cottons; the former of which is expected due to proximity, but the latter is neither expected from geographic proximity nor from likely ancestry. This unexpected inference of gene flow from Guadeloupe into Mound Key likely reflects other population processes, such as shared recent ancestry, genetic drift in small localized populations, and regional dispersal dynamics associated with cyclonic storms. Finally, the model suggests both gene flow from the Mound Key cottons into the Puerto Rican Ph site outlier (Ph_6), and from Puerto Rican PR325 population into the Guadeloupe main population. Considering their ancestry reconstructions, however, the TreeMix- suggested gene flow from Mound Key into Ph_6 may derive instead from admixture with other Puerto Rican cottons (e.g., PR_CR and PR325) that exhibit some shared ancestry with unsampled, and perhaps extinct Mound Key-like populations, whereas introgression of PR325 into GD is not supported by the structure analysis, perhaps indicating that it is an artifact of analysis in these small, isolated populations.

**Figure 3.**
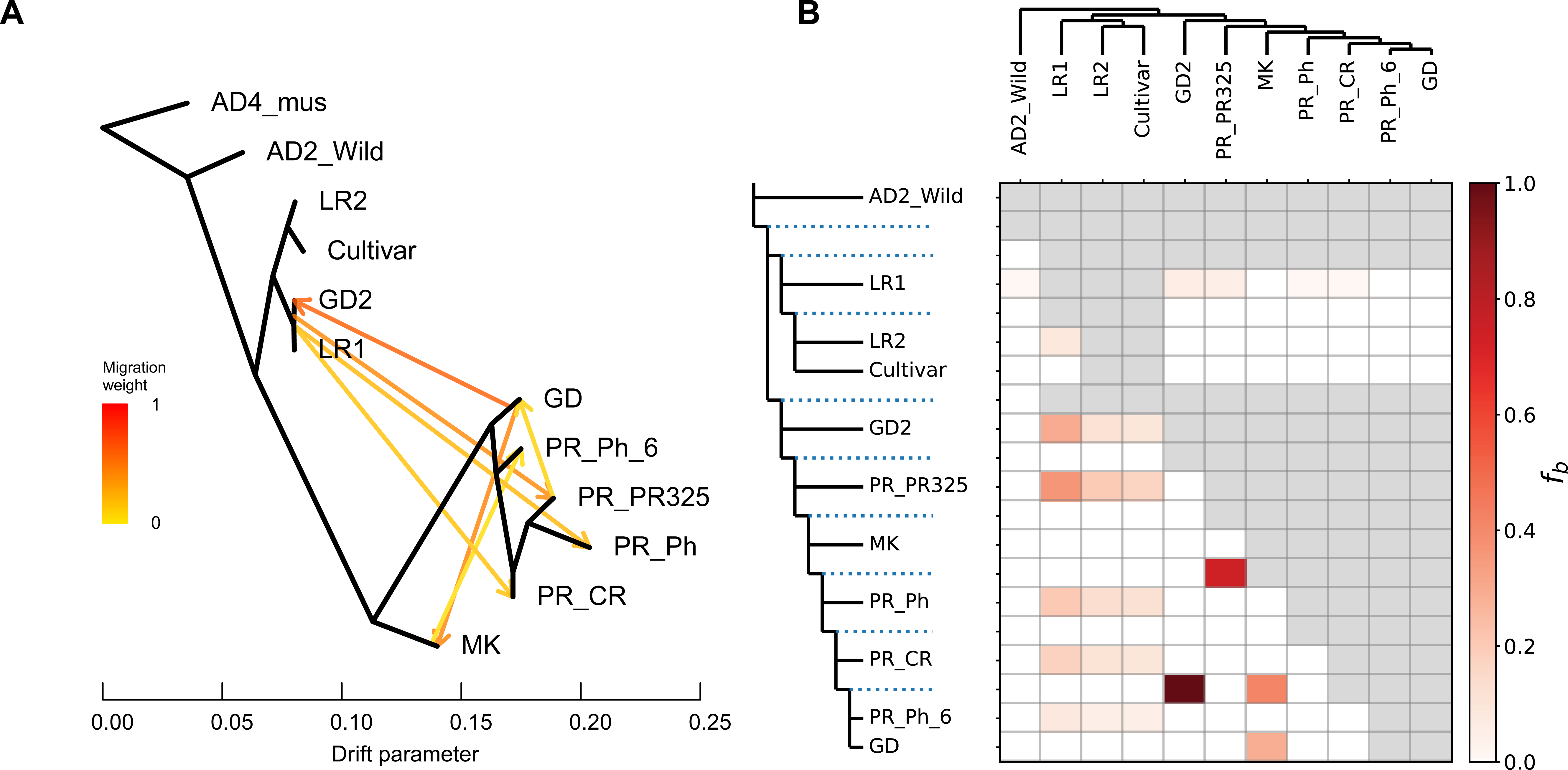
Genomic introgression tests between *Gossypium mustelinum* (AD4_mus), *G. barbadense* (AD2_wild), three domesticated cottons from germplasm collections: Landrace1 (LR1), Landrace2 (LR2), and Cultivar, and wild Caribbean and Mound Key (MK) cottons. (A) TreeMix with five migration edges shown by five branches with arrows indicating gene flow direction, and the color of the migration edges indicates migration weight. (B) Dsuite heatmap shows the level of introgression between the maximum likelihood tree nodes on the top and left. Darker color indicates values of branch introgression (*f*b).

We further analyzed the potential for genomic introgression using the ABBA-BABA test based Dsuite (Fig. 3B). The results showed that all Puerto Rico samples and the four Guadeloupe outliers (GD2) experienced different levels of gene flow from domesticated cottons, especially from Landrace1 accessions. Similar to the neighbor-joining tree and LEA structure results (Fig 2B, 2C), when assuming no gene flow, the phylogeny from TreeMix grouped the outlier PR_Ph_6 with the Guadeloupe main population (GD). The strongest introgression signal (*f*b = 1) was detected between the group containing PR_Ph_6 and Guadeloupe main population (GD) and the four geographically adjacent Guadeloupe outliers (GD2). Moreover, the ancestral node of all Caribbean wild cottons (excluding GD2) to Puerto Rican cotton PR325 also showed high introgression probabilities (*f*b = 0.7). Similar to the TreeMix result, there was also a gene flow signal detected between Mound Key and the Guadeloupe main population (GD n = 21), but given the tiny size of these islands and the great distance between them, we interpret this observation to reflect a combination of shared ancestry and population structuring (see Discussion).

### Reticulation Revealed through Plastome Phylogenomics

Chloroplast DNA is maternally inherited in *Gossypium*, and has been shown to be useful in phylogenetics and analyses of introgression (Wendel and Albert 1992; Brubaker et al. 1993; Chen et al. 2017; Wu et al. 2018; Yan et al. 2024). Among the 158 samples analyzed here, the average plastome length was 160,406 bp, similar to that previously reported (Lee et al. 2006). For phylogenetic analysis, we removed one of the large inverted repeats and all indels, resulting in a final aligned length of 134,121 bp. Maximum Parsimony (MP) (Fig. 4A; Fig. S5A) and maximum likelihood (ML) (Fig. S6A, S6B) analysis of this alignment contained 287 parsimony informative sites, only 91 of which were within the 110 cpDNA loci. This extraordinarily low level of nucleotide diversity with the chloroplast genome is not unexpected given its well-known low rate of sequence evolution, but there are few comparable datasets for this level of intraspecific sampling.

**Figure 4.**
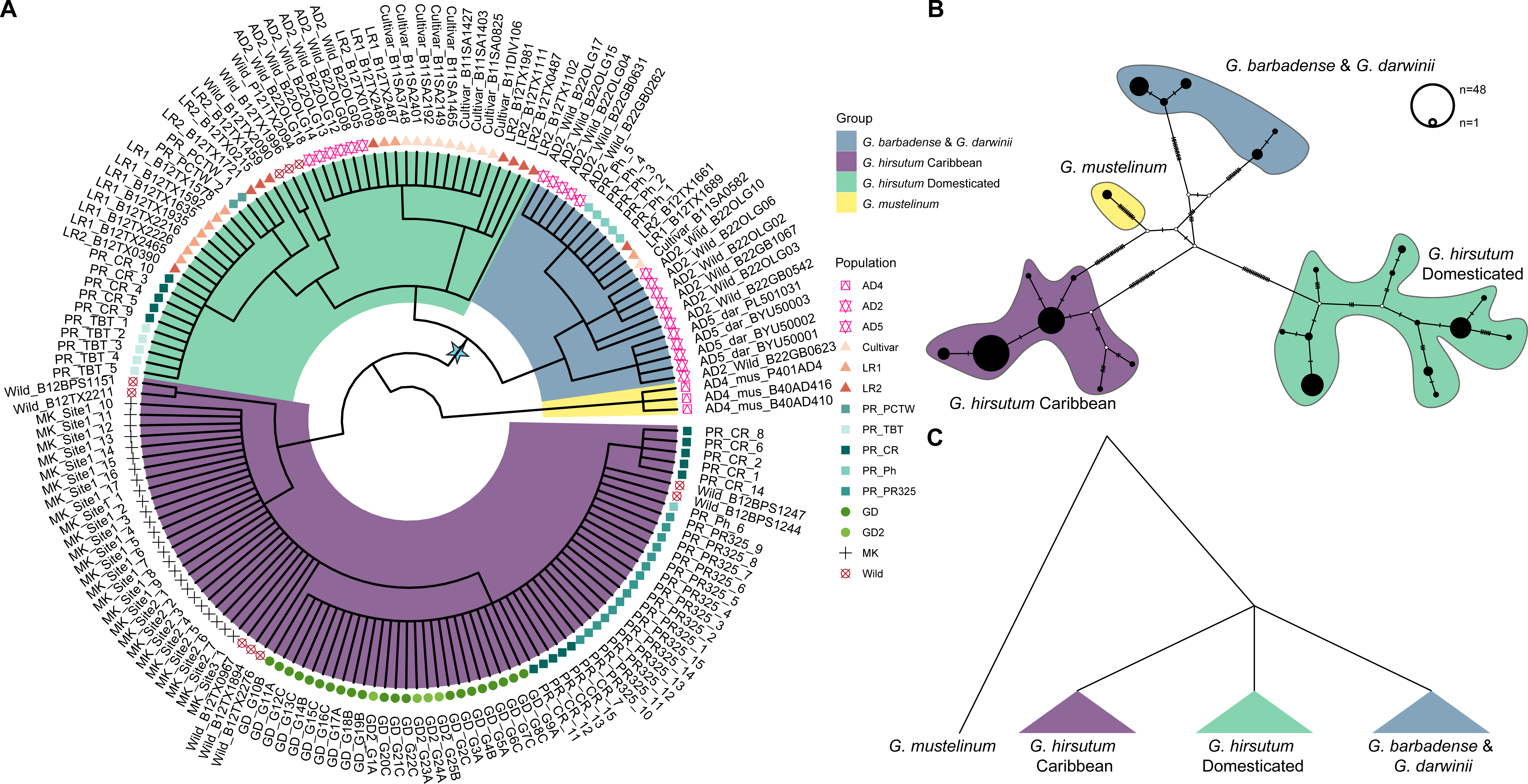
(A) Maximum parsimony phylogenetic tree for 158 *Gossypium* samples using 110 extracted gene sequences. Each tree tip is labeled by a shape and an individual ID that represents their genetic group, which was assigned using genome-wide SNPs (see Fig. 1A and Fig. S1A). Clades are colored based on assigned groups. (B) Haplotype network using a concatenation of sequences from the 110 cpDNA genes. The lines on the branch represent the nuclear changes between groups. Colors represent their cpDNA main types. (C) A cartoon tree to show genetic relationships between the three main plastome groups inferred.

Phylogenetic reconstruction using either the full genome or genes only, and MP or ML methods, both supported a similar topology and overall species relationship (Fig. 4A; Fig. S5A; Fig. S6A, S6B). Rooting with the outgroup *G. mustelinum* (AD4) (Wendel and Albert, 1992; Hu et al., 2021), the 158 samples were resolved into three major clades (Fig. 4A), namely: (1) a clade that mostly includes wild *G. barbadense* and *G. darwinii*; (2) a ‘Domesticated’ clade containing the majority of cultivated *G. hirsutum*; and (3) a ‘Caribbean’ clade, containing most of the putatively wild *G. hirsutum* collected from the Caribbean islands. Notably, however, both the ‘*G. barbadense* + *G. darwinii*’ and the ‘Domesticated *G. hirsutum*’ clades exhibited some unexpected membership and the two *G. hirsutum* clades were not monophyletic.

We further explored relationships among these plastomes using haplotype network reconstruction of both the 110 cpDNA loci and the plastomes (Fig. 4B; Fig. S5B). In both cases these networks recapitulate the three primary groups found by MP phylogenetic analysis; however, the haplotype network failed to resolve relationships among these three main groups (Fig. 4B). Importantly, although the phylogeny supported the ‘Domesticated *G. hirsutum*’ clade as sister to the two sister clades of (‘*G. barbadense* + *G. darwinii*’) clade rather than the ‘Caribbean’ clade, only two parsimony informative sites (including one homoplasious site) supported the node labelled with a star in Fig. 4A, which separates the wild *G. hirsutum* samples from the ‘*G. barbadense* and *G. darwini*’ and ‘Domesticated’ clades. Accordingly, for the purpose of this study, we treated those three groups as a polytomy (Fig. 4C). This result serves as a striking reflection of the relatively recency of species divergence and domestication when viewed on the scale of cpDNA sequence evolution.

Although the composition of each clade was largely consistent, notable exceptions were identified that may suggest historical introgression leading to chloroplast capture (Wendel and Albert 1992). While the ‘*G. barbadense + G. darwinii*’ clade contained all nine *G. barbadense* samples that we judged to be wild (see above) and all *G. darwinii* samples, it also contained *G. hirsutum* samples from the Puerto Rican population Ph (n = 5) and three domesticated *G. hirsutum* individuals previously sampled from the National Cotton germplasm collection (Yuan et al. 2021). Likewise, the ‘Domesticated’ group was largely composed of *G. hirsutum*, albeit with the unexpected inclusion of some putatively wild *G. hirsutum* and *G. barbadense* samples. As expected, the Cultivars, Landrace1, and Landrace2 *G. hirsutum* samples from Yuan et al (2021) Puerto Rico feral cottons from the TBT and PCTW sites, were all included in the ‘Domesticated’ *G. hirsutum* clade. Interestingly, however, three Wild samples from Yucatán, Mexico sequenced by Yuan et al (2021) and five individuals from the wild cotton CR site (Puerto Rico) were also included within the ‘Domesticated’ clade, as were six wild *G. barbadense* samples with mixed ancestral signals from more than one population (Fig S1C), potentially indicating a history of hybridization or introgression. All remaining wild samples, including those from Mound Key and other Caribbean islands (including seven Wild samples from germplasm collections) grouped together to form the ‘Caribbean’ clade.

### High Genetic Diversity Preserved among Caribbean Cottons

Genome-wide nucleotide diversity (π) was estimated for 121 individuals, including all four *G. hirsutum* groups from Yuan et al. (2021), Mound Key cottons, and all Caribbean wild cottons (Fig. 5A). The four outlier samples from Guadeloupe (GD2) and the single outlier sample from Puerto Rico Ph site (PR_Ph6) were excluded due to their mixed genomic signals and paraphyletic placement in the NJ tree (Fig 2B, 2C). Nucleotide diversity within populations (Fig. 5A) for the four *G. hirsutum* groups from Yuan et al. (2021) was, on average, highest for the Wild group (2.54 x 10^-3^), followed by similar levels in Landrace2 (1.95 x 10^-3^) and Landrace1 (1.90 x 10^-3^), with the lowest diversity being found in the Cultivars (9.93 x 10^-4^), as expected given the strong genetic bottlenecks accompanying cotton domestication (Wendel et al. 1992; Brubaker and Wendel 1994). Notably, however, one of the chromosomes (D08) in the cultivars exhibited elevated diversity (3.07 x 10^-3^), possibly resulting from the known structural variation in the cultivars (e.g., Hu et al. 2025).

**Figure 5.**
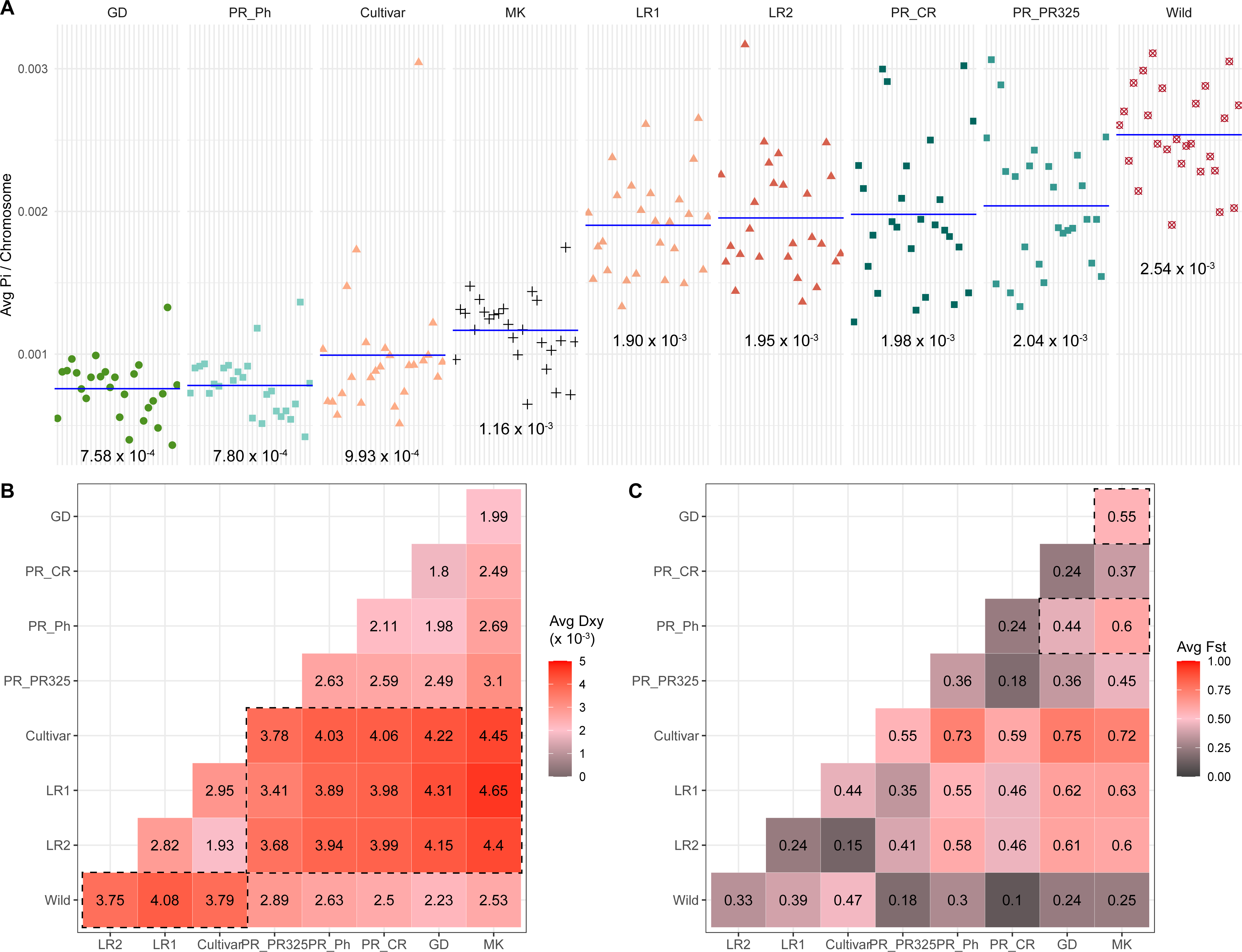
Genetic diversity and divergence measurements of four *G. hirsutum* groups and wild cottons from populations in the Caribbean and Mound Key. (A) Average nucleotide diversity (π) per chromosome. Populations/groups are distinguished by shape and color. Chromosomes are ordered along the x-axis as A01-A13, D01-D13. Mean π across all chromosomes is indicated by a blue line with values labeled below each subplot. Heatmap plot of averaged pairwise comparisons of (B) d_xy_ and (C) F_st_ between nine populations/groups. Deeper color represents higher differences between each pair, and actual values are labeled in each comparison.

Among the additional wild cotton populations analyzed here, two Puerto Rican populations PR325 (2.04 x 10^-3^) and CR (1.98 x 10^-3^) had similar diversity as in the two landrace groups, whereas Mound Key cottons (1.16 x 10^-3^) exhibited nucleotide diversities in between those of the landraces and Cultivars. Moreover, these cottons also exhibited large variation between chromosomes, which reiterated the higher divergence observed between individuals within each population (Fig 2C). In contrast, the main Guadeloupe population (GD) and cottons from the Puerto Rico Ph site (PR_Ph) showed similarly exceptionally low diversity and with lower levels of variation between chromosomes, 7.58 x 10^-4^ and 7.80 x 10^-4^, respectively, reflecting their highly inbred nature (Fig 2A; Fig. S2).

Pairwise nucleotide differences (d_xy_) were calculated between all pairs of populations (Fig. ^5^B). Compared to all three domesticated cotton groups (Landrace1, Landrace2 and Cultivars), the d_xy_ (in dash boxes) in Wild group (> 0.0038), Caribbean wild cottons (> 0.0034), and Mound Key cottons (> 0.0044) all showed sequence divergence at the upper end of the range. These data mirror the PCA results (Fig. 1B), in which the domesticated cottons and wild cottons exhibit greater sequence divergence. Among wild cotton populations, Puerto Rican site PR325 cottons exhibited higher sequence divergence (> 0.0025) from all other cotton populations, and especially when compared to Mound Key cottons (d_xy_ = 0.0031). By contrast, relatively lower divergence (d_xy_ < 0.0020) is found between the Guadeloupe (GD) and Mound Key population, and for samples from the two Puerto Rican populations at CR and Ph.

Population differentiation (F_st_) between wild and domesticated cottons showed two distinct patterns (Fig. 5C). When compared to the three domesticated cotton groups (Landrace1, Landrace2, and Cultivars), the wild cotton populations from Mound Key, Puerto Rico Ph site and Guadeloupe exhibited higher F_st_ (0.55 to 0.77), whereas the PR325 and CR site Puerto Rican cottons showed a lower F_st_ range (0.35 to 0.59). These results reiterate those from analyses of genetic structure (Fig 2C), introgression (Fig. 3A, 3B), and plastome phylogenomics (Fig. 4A), in which PR325 and CR contained more mixed ancestral signals than other wild cottons. Among the putatively wild cotton populations from Mound Key, Puerto Rico and Guadeloupe, all three pairs (in dash boxes) had F_st_ values between 0.44 to 0.60.

### Levels of Homozygosity and Heterozygosity

Long runs of homozygosity (ROH), contiguous homozygous regions inherited from both parents are informative about inbreeding and demographic history. Although ROH-based measures of inbreeding have been widely applied in humans and domesticated animals (reviewed in Shafer and Kardos 2025), their use in plants remains limited (e.g., Kumar et al. 2020; Bemmels et al. 2025). Using 26 million biallelic SNPs from 121 samples, we found the number of ROH per genome ranged from 57 to 878, with genome-wide ROH coverage varying from 1.1 % to 13.6% (Fig. 6A; Fig. S7B). Most ROH were detected in short tracts below 1 Mb (Fig. 6A), and, as might be expected, samples with higher numbers of ROH also had larger genome-wide ROH coverage (Fig. ^S^7^B^), as a metric of inbreeding coefficient based on ROH (F_ROH_). The F_ROH_ values were the highest in the Cultivars (0.092) and Landrace2 (0.081) groups, followed by the Guadeloupe main population (GD; 0.049), Landrace1 (0.043) and Wild group (0.040), with the populations from Puerto Rico exhibiting much lower homozygosity levels (0.028 Ph (n = 5); 0.023 CR, and 0.021 PR325). The lowest F_ROH_ (0.017) was found in the Mound Key population (Fig. 6C; Fig. S7A), which seems counterintuitive given its small population size and geographic isolation from other cottons.

**Figure 6.**
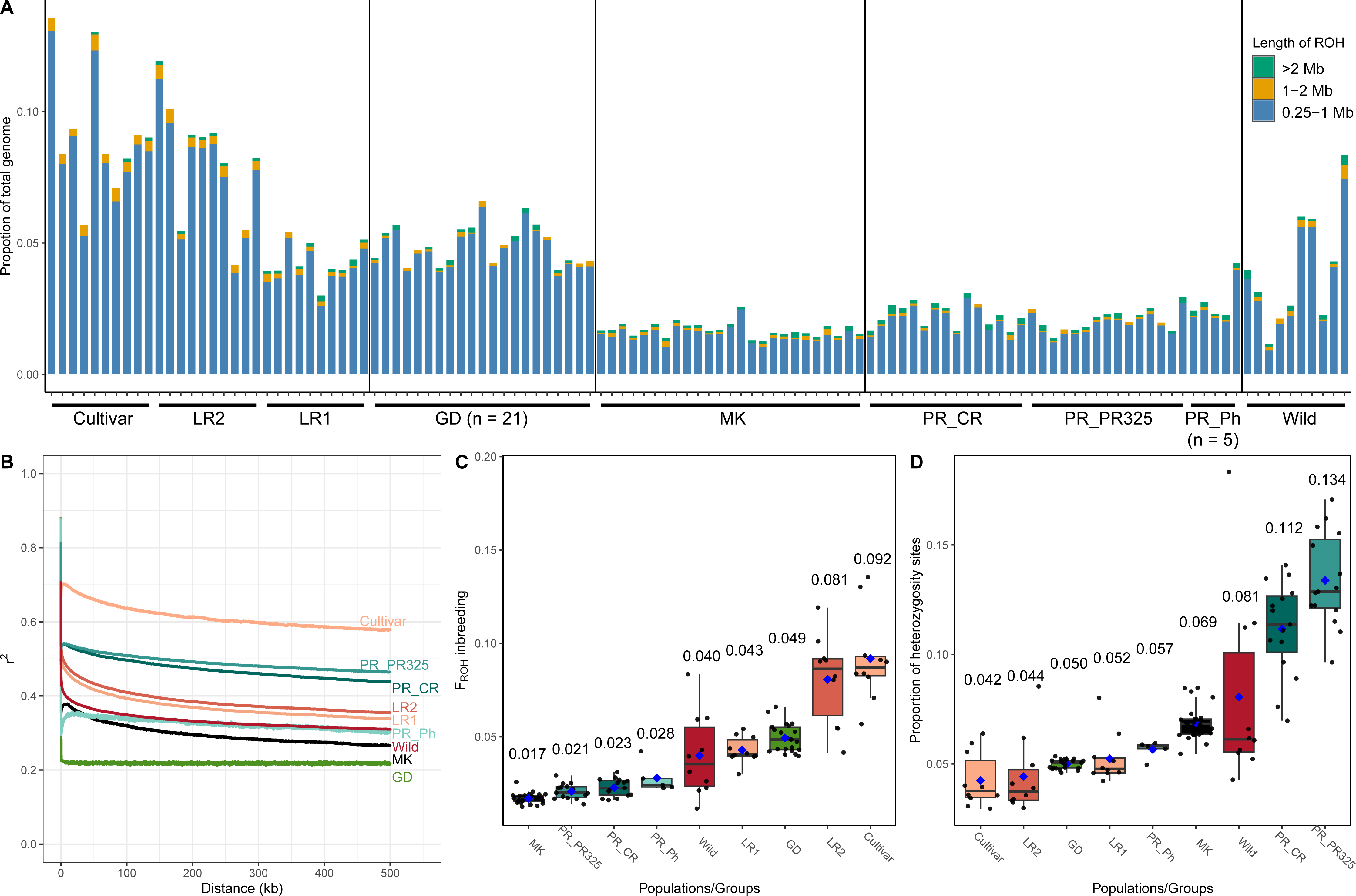
(A) Total genome proportion of runs of homozygosity (ROH) in each individual. Bar color shows the ROH in different length categories. (B) Linkage disequilibrium (LD) decay shown by the correlations of *r^2^* and the distance (kb) between SNP pairs. Each group/population is labelled by a different color. (C) F_ROH_ inbreeding coefficient and (D) Proportions of heterozygosity sites for nine populations/groups. The average F_ROH_ is shown by a blue dot and labelled above each boxplot.

Complementing the ROH analysis, we also estimated linkage disequilibrium (LD) to infer differences in historical demography that influence SNP co-occurrence (e.g., inbreeding; Fig. 6B). Previous analysis of LD decay (Yuan et al., (2021) revealed that LD remain the highest for Cultivars where pairwise SNP distances reach 500 kbp (i.e., the slowest decay), followed by faster and overlapping LD decay slopes in Landrace1 and Landrace2, with the Wild group showing the fastest decay. Among the newly sampled wild cotton populations, LD in PR325 and PR_CR decayed the slowest, falling intermediate between the landraces and the Cultivars. Conversely, Ph from Puerto Rico and the Mound Key cottons showed slower decay, exhibiting patterns similar to the Wild group. Interestingly, the Guadeloupe samples showed an initial rapid decay and then a flat *r^2^* trend which had no correlation with physical distances of SNPs.

As a final measure of genome-wide diversity, we calculated the proportion of heterozygous sites in each genome. The highest value was observed for the two Puerto Rican populations with a history of admixture, i.e., PR325 (13.4%) and CR (11.2%), where heterozygosity was even higher than the Wild cotton group (8.1%) (Fig. 6D). In contrast, the Ph main population from Puerto Rico showed only half this amount of genome-wide heterozygosity (5.7%) relative to PR325 or PR_CR. The Mound Key cottons (6.9%) exhibited larger proportions of heterozygous sites than the Ph population. As expected, given the strong selection and inbreeding accompanying true-breeding line development in cotton cultivars, the domesticated cottons had the lowest overall heterozygosity (Fig. 6B, 6C). In particular, the Cultivar group had the lowest heterozygosity (4.2%) compared to Landrace2 (4.4%) and Landrace1 (5.2%). Notably, the Guadeloupe main population (GD), despite being a wild cotton population, showed a similarly low proportion of heterozygous sites (5%) as the domesticated cotton group Landrace1. This may reflect a small effective population size as well as a protracted history of localized inbreeding.

### Understanding Divergence of Wild Cottons under Coalescent model

To investigate the evolutionary history of wild *G. hirsutum* populations, we first used Tajima’s *D* as a proxy to assess deviations of rare allele distributions from neutral allele frequency distributions (Nielsen 2001). Given the known sensitivity of this statistic to sample size (Subramanian 2016), the Puerto Rican Ph cotton population (n=5) was removed from the analysis. Tajima’s *D* (Fig. 7A) was negative for Mound Key (-0.444) and slightly negative for Puerto Rico CR cottons (-0.146), indicating an excess of rare alleles compared to the neutral model, usually reflecting recent population expansion or purifying selection. Conversely, Tajima’s *D* was positive for both the Guadeloupe main population (GD; 0.949) and Puerto Rican PR_PR325 (0.439), indicating shortage of rare alleles and excess of intermediate-frequency alleles, often a signature of past population contraction or balancing selection.

**Figure 7.**
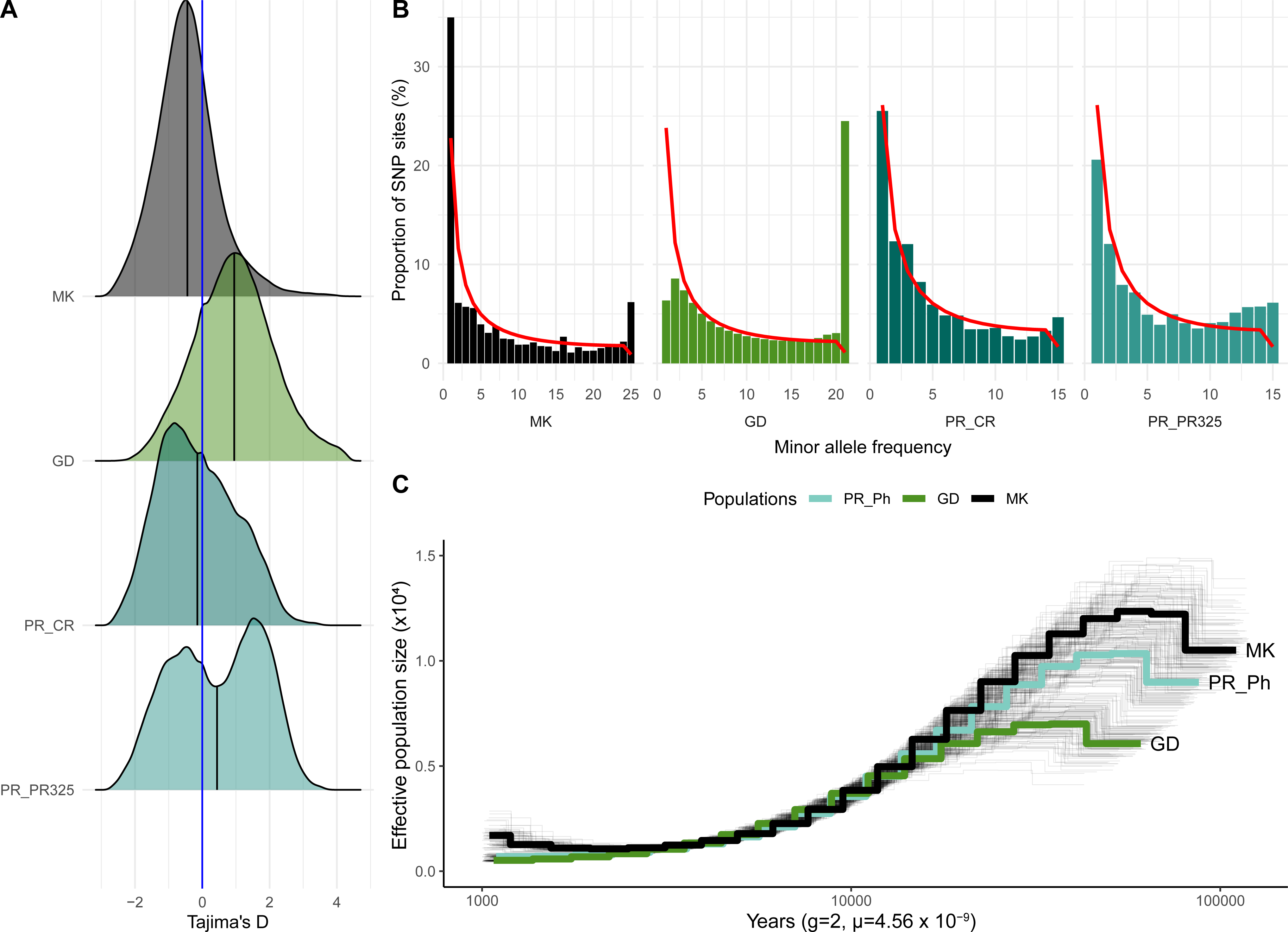
A) Density plot of Tajima’s *D*. B) The folded minor allele frequency plot against the proportions of the total variant sites for four populations, with the expected variant site distribution along different minor allele frequencies plotted as a red line. C) PSMC analysis of inferred effective population size change (y-axis) over evolutionary time (x-axis), where each of the 100 replicates for each individual shown is shown as the thin black line and the averaged *Ne* changes across all replicates for each individual is colored and labelled.

As it is well-understood that selection, admixture, population history, subpopulation structure, and nonrandom sampling could affect Tajima’s *D* estimation (Nielsen 2001; Ramos-Onsins and Rozas 2002; Moeller et al. 2007), we further explored the observed and expected site frequency spectrum (SFS) to reveal allele frequencies and distributions in four populations, where deviations from the expected allele distribution can also provide insight into selection or changes in population size (Pavlidis and Alachiotis 2017; Beichman et al. 2018). Since the ancestral allele is unknown, we calculated the observed folded SFS for each population based on the minor allele frequency (MAF). Notably, Guadeloupe cotton only showed one-third to one-fourth of variable sites (4.7 M) when compared to Mound Key cottons (15.3 M), Puerto Rican cottons PR325 (17.5 M) and CR (20.1 M) (Fig. S8A), further underscoring its general uniformity.

When compared to the expected SFS (calculated using Watterson’s theta based neutral modelling (Hudson 2015)), all four wild cotton populations exhibited an excess of common alleles (i.e., the last peak of the SFS plots in Fig. 7B; Fig. S8A), although these were proportionately much higher in the Guadeloupe main population (GD). Guadeloupe (GD) exhibited both a deficit of rare minor alleles (i.e., first peak in SFS) and contained a large peak representing common alleles (minor allele frequency = 0.5), results congruent with a positive Tajima’s *D* and suggestive of a recent population contraction (bottleneck). Both observations are also congruent with the aforementioned low diversity and high inbreeding, all of which indicate a general uniformity among the members of this population. Conversely, the Mound Key population exhibited an excess of rare variants, which, combined with a negative Tajima’s *D*, may indicate a history of recent population expansion or gene flow. In contrast, two Puerto Rican cotton populations PR325 and CR both exhibited excess of intermediate to high frequency alleles, and deficit of rare alleles in PR325, which may suggest recent admixture followed by inbreeding in both populations.

Finally, we also applied Pairwise Sequentially Markovian Coalescent (PSMC) modeling to to infer long-term effective population size (*Ne*) changes, acknowledging that the strong population structure from inbreeding, admixture between populations, and short divergence time may violate model assumptions and likely reduce our resolution or confound our results (Wang et al. 2016; Bansal and Nichols 2025). Indeed, PSMC analysis recovered two distinct patterns (Fig. S8B): one in which populations exhibited unexpected historical growth and one where they consistently declined. The first pattern was observed in the Puerto Rican cotton populations CR and PR325, both of which had excess heterozygosities due to introgression (Fig. 6D; Fig. 3A, 3B), reducing reliability of its effective population size (*Ne*) estimates (Fig. S8B). By contrast, the Mound Key, Puerto Rico Ph, and Guadeloupe (GD) populations exhibited generally low *Ne*, estimated at less than 20,000 individuals 100,000 years ago (ya), then decreasing to nearly 0 (Fig. 7C), the latter is consistent with the small number of individuals observed at each site and the human-induced population crash associated with habitat destruction and range restriction. This trend, however, could also reflect subpopulation structure or other violations of the model (Bansal and Nichols 2025) and should be interpreted with caution.

## DISCUSSION

Plant domestication represents one of the most transformative shifts in human history, enabling the development of agriculture and the rise of complex societies (Meyer and Purugganan 2013). Through selective propagation, wild species were genetically altered to produce more uniform, desirable phenotypes that comprise the elite cultivars grown today. An unintended and unfortunate consequence of this selection process, however, has been the winnowing of genetic diversity, potentially limiting the ability of modern crops to adaptively respond to pathogens, pests, and environmental stressors. Accordingly, the genetic resources represented by wild crop relatives are considered a valuable resource for introducing traits and genetic variability into modern inbred crops (Hajjar and Hodgkin 2007; Warschefsky et al. 2014; Brozynska et al. 2016; Zhang et al. 2017). For many crop species, however, ancestral wild gene pools have been depleted by habitat destruction or other human-mediated impacts, and/or face extinction (Ford-Lloyd et al., 2011; Maxted et al., 2010).

Given the vital role that *G. hirsutum* plays in global commerce, it is remarkable that so little is understood about the species as a wild plant. Prior to human disturbance and domestication, in natural settings wild cotton occurred as a fully integrated woody perennial shrub to small tree in the arid to semiarid scrub forests that characterize some of the leeward coastal habitats in a broad swath of the Caribbean, ranging from perhaps Trinidad and Tobago in the south to as far north as Tampa Bay, Florida, with the best developed population systems in the northern part of the Yucatan Peninsula (Stephens and Phillips 1972; Fryxell 1979; Wendel et al. 1992; d’Eeckenbrugge and Lacape 2014; Hu et al. 2021). Although this aggregate geographical range is extensive, actual populations typically are widely scattered and may be highly inbred. Complicating our understanding of the status and diversity of wild populations has been the effects of human history, which includes both widespread cultivation of cotton in agricultural settings and habitat destruction of the native vegetation in which wild cotton occurs, leading to further habitat fragmentation. Finally, cultivated cottons are interfertile with wild forms, thus leading to gene flow into natural settings and the establishment of feral derivatives in native vegetation. Due to these many complexities, understanding the domestication genetics and genetic diversity of truly wild *G. hirsutum* has been challenging. A corollary is that there is little understanding of the actual proportion and source of wild diversity that has been captured or represented in modern elite gene pools.

Here we use a previously generated phylogenomic framework (Yuan et al. 2021) to evaluate the genome-wide diversity and population structure in newly collected wild and putatively wild populations of *G. hirsutum* from the Caribbean. Analysis of field-collected cotton from multiple sites in Puerto Rico and Guadeloupe reveals novel pockets of cotton genomic diversity not yet described, which both expand our understanding of wild diversity in the species and form a potential resource for targeted breeding. Using population genomics, we quantify diversity, divergence, and possible gene flow dynamics among different island populations, as well as introgression from a second domesticated polyploid cotton species found in the Caribbean, *G. barbadense*. Our results show that populations of Caribbean *G. hirsutum* consists of three broad groups, purely wild *G. hirsutum*, feral derivatives of previous stages in the domestication continuum, and plants derived from ancient hybridization and introgression from *G. barbadense*. Below we discuss the origins and diversity of Caribbean wild cottons.

### Genomic Diversity among Domesticated Cottons

A recent phylogenomic context for understanding *G. hirsutum* diversity was generated by Yuan et al (2021) using mostly germplasm from the Germplasm Resources Information Network (GRIN; (Byrne et al. 2018)). This analysis grouped accessions into four broadly encompassing genetic groups, one comprising obsolete as well as modern cultivars, and three others representing collections from the wild and indigenous (pre-Columbian) domesticated range. These three groups, each heterogeneous, were termed Landrace1, Landrace2, and Wild, with Landraces 1 and 2 also aligning loosely with geography, the former broadly encompassing the Caribbean and the latter consisting of early domesticated forms from Central America. The utility of this framework was recently demonstrated by using it to verify that cotton populations from Mound Key, Florida are truly wild as opposed to feral (Ning et al. 2024) and to demonstrate that this population represented a novel or previously unknown pocket of genetic diversity.

Here we extend this analysis to demonstrate the utility of this framework to distinguish other “wild-looking” feral derivatives from truly wild cottons. These feral plant populations are highly heterogeneous and are thought to have been derived from primitively domesticated forms that repeatedly escaped from human into natural settings over the past couple of millenia, as cotton cultivation spread from its ancestral home to encompass much of the drier American subtropics and tropics, up to and including more recent escapes and introgressants (from agricultural fields) in the post-colonial era from larger-scale cotton field agriculture. Using representatives from the framework provided by Yuan et al (2021), we found that all samples from two sites in Puerto Rican (PCTW and TBT) are genomically similar to the domesticated and landrace cottons, suggesting that they represent feral cottons or early-domesticated forms (Fig. 1C) rather than truly wild relict. This conclusion was further supported by plastome variation, which revealed that cottons from these two sites contain the same cpDNA type as the domesticated cottons (the ‘Domesticated’ group in Fig, 4A), which was distinct from the wild cottons. Notably, the morphology of these plants was ambiguous, in that they form multi-branched shrubs, consistent with wild cotton plant architecture, but with seed trichomes (“fibers”) that indicate the influence of domesticated cotton, in that they were white and longer than in truly wild forms. This quasi-wild phenotype is, in fact, quite common, as it has been noted that the feral cotton may regain wild cotton morphology in architecture after escaping cultivation and becoming reestablished in natural or disturbed sites (Hutchinson 1951; Fryxell 1979; Brubaker and Wendel 1994; d’Eeckenbrugge and Lacape 2014).

Similar to other domesticated crops (reviewed in Zhang et al. 2017), domestication led to a series of genetic diversity reductions from wild cottons to early domesticated cottons (i.e., landraces), and eventually to the modern cultivars (Fig. 5A). Despite this decline, domesticated cotton gene pools contain a substantial proportion of novel genetic variations that were absent in wild cottons (Ning et al. 2024). Principal component analysis and neighbor-joining trees support a close genetic relationship, placing the domesticated cottons as sister to wild cottons (Fig. 1B, 1C). These findings highlight the reciprocal introgression history between *G. hirsutum* and *G. barbadense*, with all domesticated *G. hirsutum* containing different levels of genomic introgression signals from *G. barbadense* (Yuan et al. 2021). Although nuclear gene flow was not directly detected in this study, plastome variation provided additional genetic introgression evidence (Fig. 4A), in which the ‘Domesticated’ group contained six *G. barbadense* samples that had more mixing signals in nuclear genetic structure analysis (Fig S1C), and the ‘*G. barbadense* & *G. darwinii*’ group had three domesticated *G. hirsutum* individuals from Landraces and Cultivars. The reciprocal introgression likely contributed to increased genomic variation in cultivated *G. hirsutum* and resulted in higher divergence levels compared to the wild relatives.

### Origins of Semi-Wild Cottons

When the feral derivatives are excluded, all remaining wild-collected cotton samples included in this study grouped with the Wild germplasm group from Yuan et al (2021) (Fig. 1B, 1C), using both PCA and phylogenetics. This result notwithstanding, population structure analysis revealed that some of these have signals of mixed ancestry. That is, unlike the general uniformity exhibited by the Mound Key cottons and most of the cottons from Guadeloupe and Puerto Rico site Ph, cotton collected from the Puerto Rican sites PR325 and CR exhibited patterns of genetic variation that implicate an origin or influence from more than one population, as did the small population from Guadeloupe (GD2). Specifically, these “semi-wild” cottons contained different levels of ancestral signals from a proximal wild cotton population along with mixed signals from various other sources, including the domesticated pool (Fig. 2C). Genomic introgression tests (see below) highlighted Landrace1 (and related groups) as a possible source of introgression (Fig. 3A, 3B), congruent with the localization of that group in the Caribbean (Yuan et al. 2021). Notably, the morphology of these semi-wild cottons was consistent with the wild cotton phenotype, e.g., multi-branched architecture and seeds with short-brownish fibers, providing morphological evidence of historical genomic introgression.

Further analysis of these samples revealed varying degrees of “contamination” from domesticated sources. In Guadeloupe, for example, the presence of domesticated alleles among the four semi-wild samples (i.e., GD2) was so pervasive that TreeMix grouped GD2 with Landrace1, suggesting that the direction of introgression was from the wild (main) GD population into a Landrace1-like background (Fig. 3A). Notably, when collecting these, Georges Ano noted (J. Wendel, pers. comm.) that these four cottons appeared as “hybride Marie Galante - Yucatenense”, the former representing a semi-domesticated race of *G. hirsutum* known from (and named for) the nearby island of Marie-Galante (Ano et al. 1983) and the latter representing the truly wild form known as *G. hirsutum* race Yucatanense (Hutchinson, 1951; Fryxell, 1979; Wendel et al., 1992). The presence of both “primitive” (i.e., race Yucatanense) *G. hirsutum* and *G. hirsutum* race Marie-Galante on Guadeloupe (Ano et al. 1982), along with a history of cotton cultivation during the colonial period (Hoy 1962), offers an explanation for the introduction of domesticated alleles into a predominantly wild background. Given our observations and the history of both wild and early domesticated forms on Guadeloupe, it is reasonable to suggest that these semi-wild cottons originate from gene flow between the wild cottons that inhabit Guadeloupe and one (or more) feral forms. Moreover, the four semi-wild outliers in Guadeloupe (GD2) form two divergent lineages (GD2_G25B with GD2_G1A, and GD2_G24A with GD2_G23A) in the neighbor-joining tree (Fig. 2B), perhaps reflecting historical differences in genomic recombination and segregation (Fig. 2C). Perhaps more striking is the observation that these four samples (collected within meters of one another) were growing at a site that is closely adjacent to the Guadeloupe main population (GD), yet no signature of admixture with domesticated cottons was evident in the GD population.

A similar interpretation applies to the semi-wild cottons found on Puerto Rico, which also are interpreted here as having mixed ancestry arising when natural, wild populations of *G. hirsutum* encountered feral remnants of previously cultivated material. Among the Puerto Rican sites, both CR and PR325 were composed solely of cottons with varying proportions of mixed ancestry (Fig. 2A, 2B). The PR325 population exhibited strong introgression signals from Landrace1 (Fig. 3B), which supports hybrid ancestry for the PR325 cottons. By contrast, the CR population also exhibited some (lesser) evidence of introgression from Landrace1; however, it also exhibited evidence of plastome introgression that was absent in the PR325 cottons. Specifically, three different plastome types were found among the CR cottons (Fig. 4A), including five individuals with the ‘Domesticated’ type , and the remaining ten individuals containing the ‘Caribbean’ type (Fig. 4). Therefore, while the semi-wild cottons from the Puerto Rican sites CR and PR325 both likely result from introgression with feral cottons, the timing and contemporary outcomes of this contact appear variable between sites.

### Genomic Diversity of Semi-Wild Cottons

Gene flow from the domesticated crops to their wild progenitors has been documented in other crop plants (reviewed in Gepts 2014), such as common bean (Papa and Gepts 2003), maize (Hufford et al. 2013), barley (Russell et al. 2011), rice (Chen et al. 2004; Jin et al. 2018), and wheat (Nyine et al. 2020). These introgression events resulted in altering the genomic landscape and genomic diversity of wild crop relatives by introducing the foreign domesticated gene pools. Similarly, although the Puerto Rican CR and PR325 cottons had elevated heterozygosity and nucleotide variations relative to other wild cotton populations (consistent with gene flow; Fig. 5A; Fig. 6B), these two semi-wild cotton populations also exhibited similar long runs of homozygosity as inbred wild cotton populations and elevated LD, the latter of which exceeds both landraces (Fig. 6D). These seemingly contradictory observations reveal another dimension to the genomic history of the Puerto Rican cottons, i.e., the presence of feral *G. barbadense* on the island. Cotton cultivation in Puerto Rico predates Christopher Columbus; however, it was not until the late 18th century when large-scale production commenced (Rodríguez et al. 1956). By the early 20th century, *G. barbadense*, a South American native, was in broadscale cultivation while the relictual populations of wild *G. hirsutum* faced targeted eradication (Stephens 1976); however, introgression between the two species was also employed in developing new cotton varieties (Stephens 1975; Percy 2009; Viot and Wendel 2023). Previous studies have noted that the history of genomic introgression from *G. barbadense* into Landrace1 (Yuan et al. 2021) remains evident, while others note the propensity for hybrid breakdown in crosses between *G. hirsutum* and *G. barbadense* (Hu et al. 2019). With these considerations, it is tempting to suggest that the high genetic nucleotide variation and slower LD decay observed in the cottons from PR_CR and PR_PR325 may reflect the remains of additional historical admixture from genetically incompatible parents (e.g., Nyine et al. 2020); that is, these cottons may result from relictual populations of wild *G. hirsutum* encountering feral cottons that contained genomic segments of *G. barbadense*. This complex ancestry is grounded in the history of the island, and accounts for the observations of increased genetic diversity and long LD. The slower LD decay trend in these two populations may have been attributed to reduced efficiency of recombination in regions of interspecific introgression; however, this needs further studies.

### Divergence among Wild Cottons

Understanding the genetic diversity among crop wild relatives has been a central goal of modern crop breeding (Bohra et al. 2022). However, only limited plant crop wild relatives have been studied for levels and patterns of genomic diversity using whole genome level data across multiple wild populations or landraces (e.g., wheat (Cheng et al. 2024) and maize (Hufford et al. 2012)). Complementing previous efforts to understand wild cotton diversity using different tools and sampling regimes (Wendel et al. 1992; d’Eeckenbrugge and Lacape 2014; Yuan et al. 2021; Ning et al. 2024), the present study reveals hidden genetic diversity within the Caribbean islands Guadeloupe (GD, n = 21) and Puerto Rico (Ph site). These populations, like the Mound Key cottons (Ning et al. 2024), are found exclusively in arid to seasonally arid coastal regions, where they form small, inbred populations spanning just a few hundred meters along the coastline. Although all three populations had similar recent reductions in *Ne* (Fig. 7C) and generally low genetic nucleotide diversity (Fig. 5A), each population was characterized by its own unique demographic history.

Florida Mound Key cottons, which consists of two subpopulations (Fig. 2C) as previously reported (Ning et al. 2024), showed high population divergence from both the Caribbean wild cottons (Ph) and Guadeloupe (GD), with F_st_ 0.60 and 0.55 (Fig. 5D), respectively. Mound Key cotton also showed a diverged plastome type sister to all Caribbean cottons (Fig. 4A). In addition, Mound Key cottons had the largest amount of excess minor rare alleles and a negative Tajima’s *D* relative to equilibrium expectation (Fig. 7A, 7B), which may indicate the population expansion of Mound Key cotton from initial seed dispersal to the island and population establishment. Perhaps Mound Key cottons belong more generally to a regionally genetically more coherent set of wild cotton populations from southernmost Florida, with the Mound Key cottons retaining different portions of the broader ancestral polymorphism, and therefore appearing as separate populations.

Populations at the Puerto Rico Ph site and Guadeloupe (GD) both contained only one inbred population with lower F_st_ (0.44) when compared to Mound Key cotton (Fig. 5D). These two populations showed similar levels of nucleotide diversity and heterozygosity (Fig. 5A; Fig. 6D), the Guadeloupe main population was particularly characterized by its genome homogeneity. Although Guadeloupe cotton contained considerably larger heterozygosity sites (5%) than Cultivars or Landrace2 (Fig. 6D), these sites were likely all fixed in the population and resulted in a right shift of peak in SFS (Fig. 7B). The fixed heterozygous sites and lack of variation between individuals in Guadeloupe may also have led to insufficient genetic diversity to differentiate LD at short or long distances and, therefore, create a flat LD decay trend (Fig. 6C). In addition, lack of rare alleles in Guadeloupe also resulted in a positive Tajima’s *D* > 0, suggesting a sudden contraction in population size.

In contrast, all five individuals of the Ph population had a plastome variation that was grouped with wild *G. barbadense* in the ‘AD2_AD5’ group (Fig. 4A). Given the uniform ancestry in the Ph main population (Fig. 2C), this might indicate the ancient introgression from *G. barbadense* in the Ph site populations, or a recent introgression with strong impact from genetic drift (Bemmels et al. 2025). An additional Puerto Rican outlier (PR_Ph_6) similarly exhibited mixed ancestry, instead of grouping with all other five samples from the same Puerto Rico Ph site, showed over 50% mix of ancestral signals from both Guadeloupe main population (GD) and PR_CR (Fig. 2C).

Different from all other Ph individuals that showed a plastome type nested in the ‘*G. barbadense* + *G. darwinii*’ group (Fig. 4A), the outlier Ph_6 plastome was identical to all Guadeloupe and PR325 types, indicating possible gene flow from different islands. Given the limited sample size, this may also reflect additional gene flow resulting from human activities or natural dispersal events.

### Conservation Insights

As a biodiversity hotspot (Maunder et al. 2008), the Caribbean region, with more than 700 islands, provides a vast yet highly fragmented habitat, fostering the dispersal and diversification of wild cottons. The diverged microhabitats in each island and large effects from drift as well as geographic isolation provide new opportunities for novel adaptive alleles to evolve in each small relictual population. As a result, many discrete populations with unique genetic content may remain unrecognized among the islands.

Echoing the previous ecological niche modeling results (d’Eeckenbrugge and Lacape 2014), the distribution of semi-wild or wild cottons at Puerto Rico and Guadeloupe exhibited a narrow range at two islands (Fig. 1A) and in drier coastal regions (e.g., Fitzpatrick and Giovas 2021). Puerto Rican cottons were collected at multiple sites, and it appears that a few of these are arguably truly wild. In addition, the wild cottons from Guadeloupe described here were collected from one small region across perhaps several hundred meters, but these were almost identical siblings. These rare and naturally uncommon wild cottons indicate their vulnerability to climate change-induced perturbations from rising sea levels and storms, and/or habitat eradication from human activities. In addition, with documented precolonial human activities that started 6000 to 7000 years ago (Fitzpatrick and Keegan 2007), and the fact that cotton farming was at one time a dominant component of the agriculture in both islands (Viot and Wendel 2023), wild cottons experienced and may continue to face genomic contamination from feral cotton derivatives, such as the semi-wild cottons mentioned in this study.

One can easily envision that prior to human-mediated disturbance, wild cottons were once more broadly distributed in the drier coastal regions of the Caribbean basin, as shown by ecological niche modeling (d’Eeckenbrugge and Lacape 2014). Populations most likely became established by long-distance dispersal and were eliminated by seasonal storms and hurricanes, which together generated an aggregately large and shifting range for the species but one in which populations were often highly isolated and small in size. Thus, the present relative rarity further constrains natural patterns of diversity, with reproductive isolation between demes increasing generalized inbreeding and genetic depauperatization. This dynamic, quantified to a certain extent in the present study, serves to justify the need for greater collection, evaluation, and germplasm preservation of this valuable cultivated cotton species.

## Acknowledgments

We thank our collaborator and long-time cotton aficionado Georges Ano for collecting the Guadeloupe cotton samples. We thank Iowa State University Research IT for technical support; Kathleen Foster-Wendel for assistance with field collections; and Matthew Hufford and Xinlin Yuan for discussion. We also thank Kenneth McCabe and Anna Tuchin for assistance with plant care. We especially thank Emma Miller for growing the Guadeloupe plants, an experience she assures us was as unpleasant as it was botanically productive.

## Author contributions

J.F.W. conceived the project, secured the funding and collected the samples. W.N. co-designed the experimental analysis, wrote the manuscript and created all figures, with supervision from co-last authors C.E.G. and J.F.W.. W.N., C.E.G., G.H., and D.Y. contributed to data analysis. M.A.A and J.A.U contributed to reference genome assembly and annotation. Y.D., C.H., Z.V.M, and O.P. extracted DNA and generated genomic libraries. C.H., M.A.A., and D.G.P. contributed to genome sequencing.

## Data availability

Bioinformatic pipelines are uploaded to: https://github.com/Wendellab/CaribbeanAD1; All sequence data generated in this study are available: Guadeloupe cotton (PRJNA603025) and Puerto Rico cotton (PRJNA1226603).

