## Supplementary figures and images for "Comparative population genomics of relictual Caribbean island *Gossypium hirsutum*"

### Fig. S1

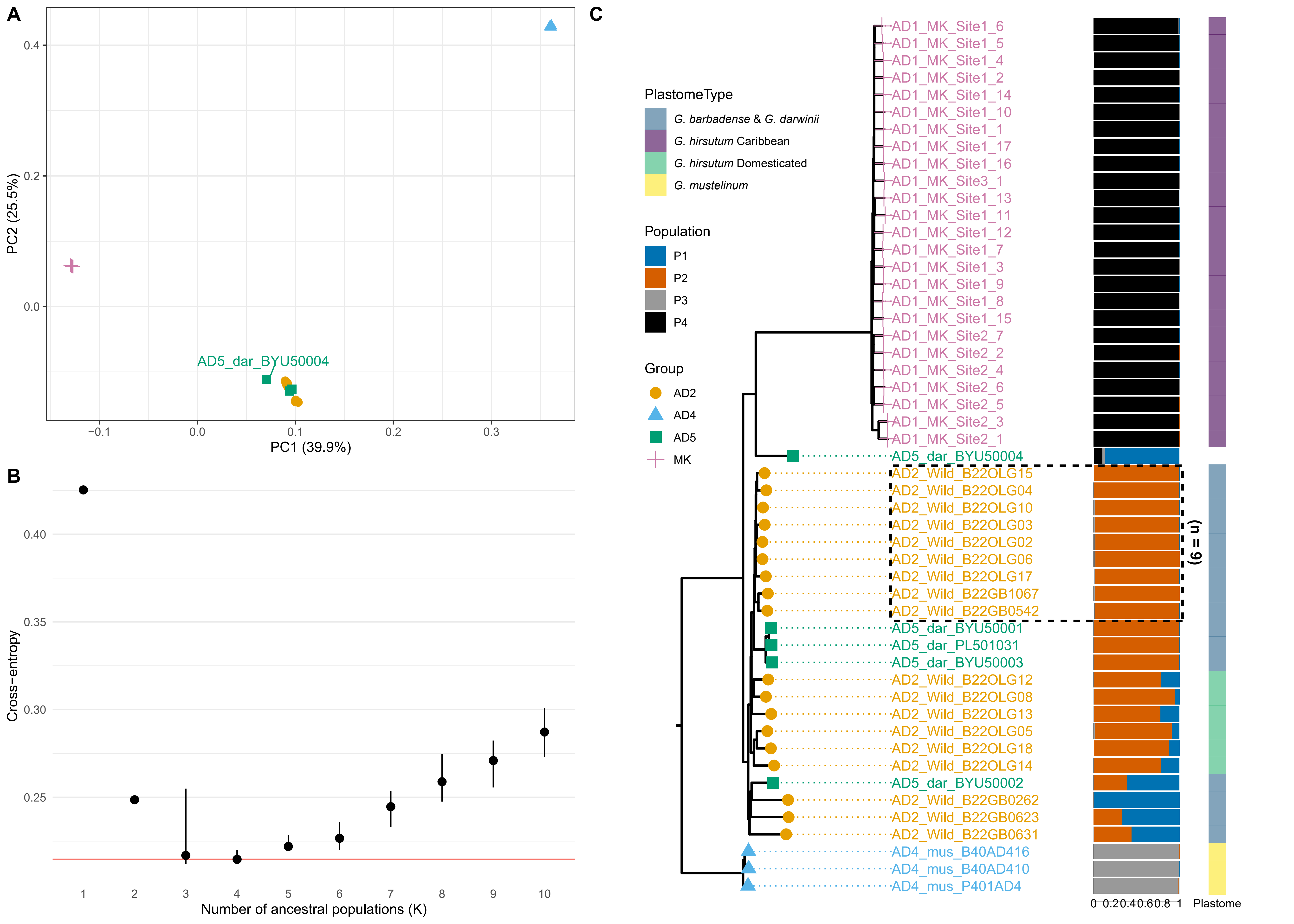

### Fig. S2

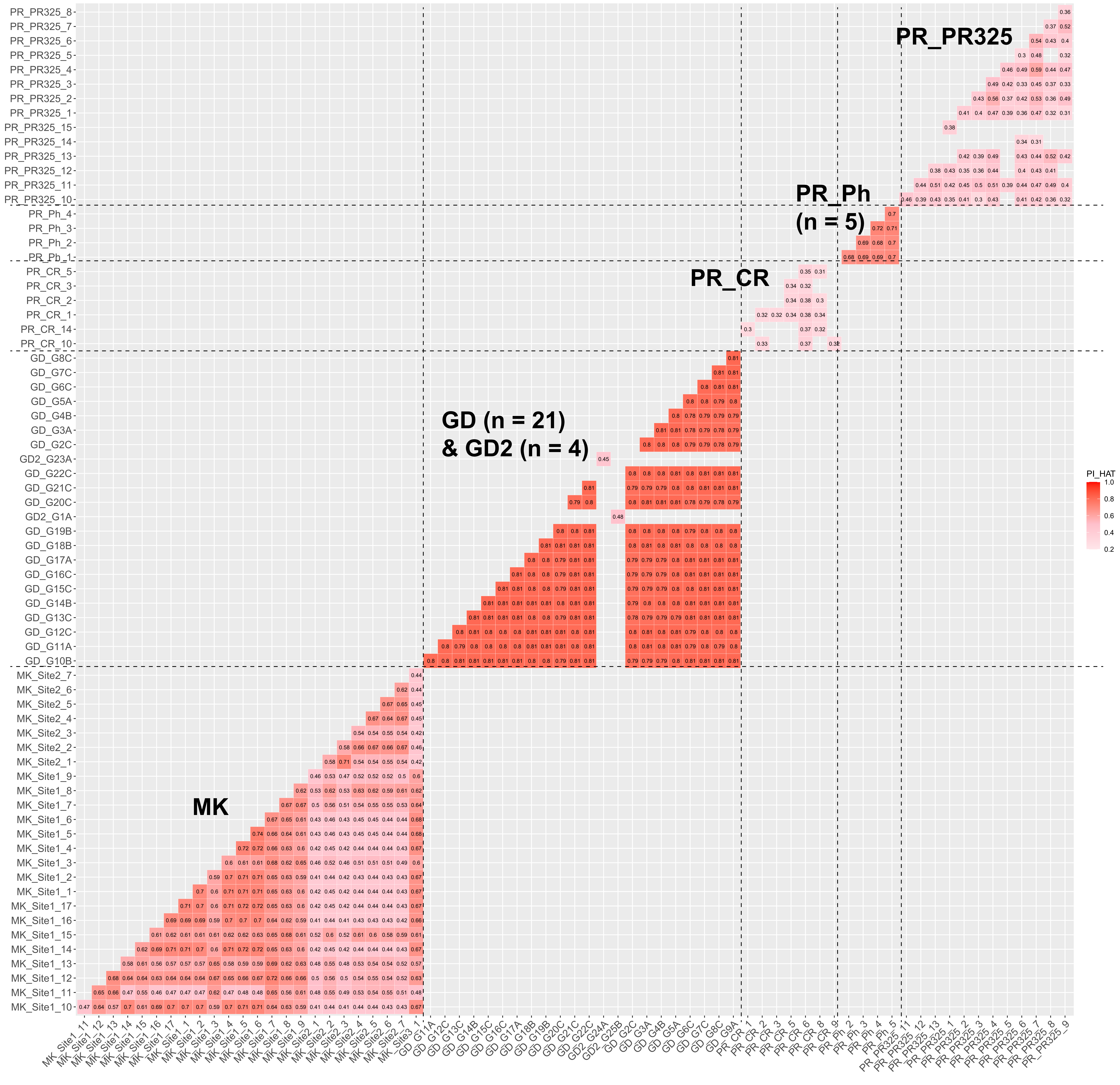

### Fig. S3

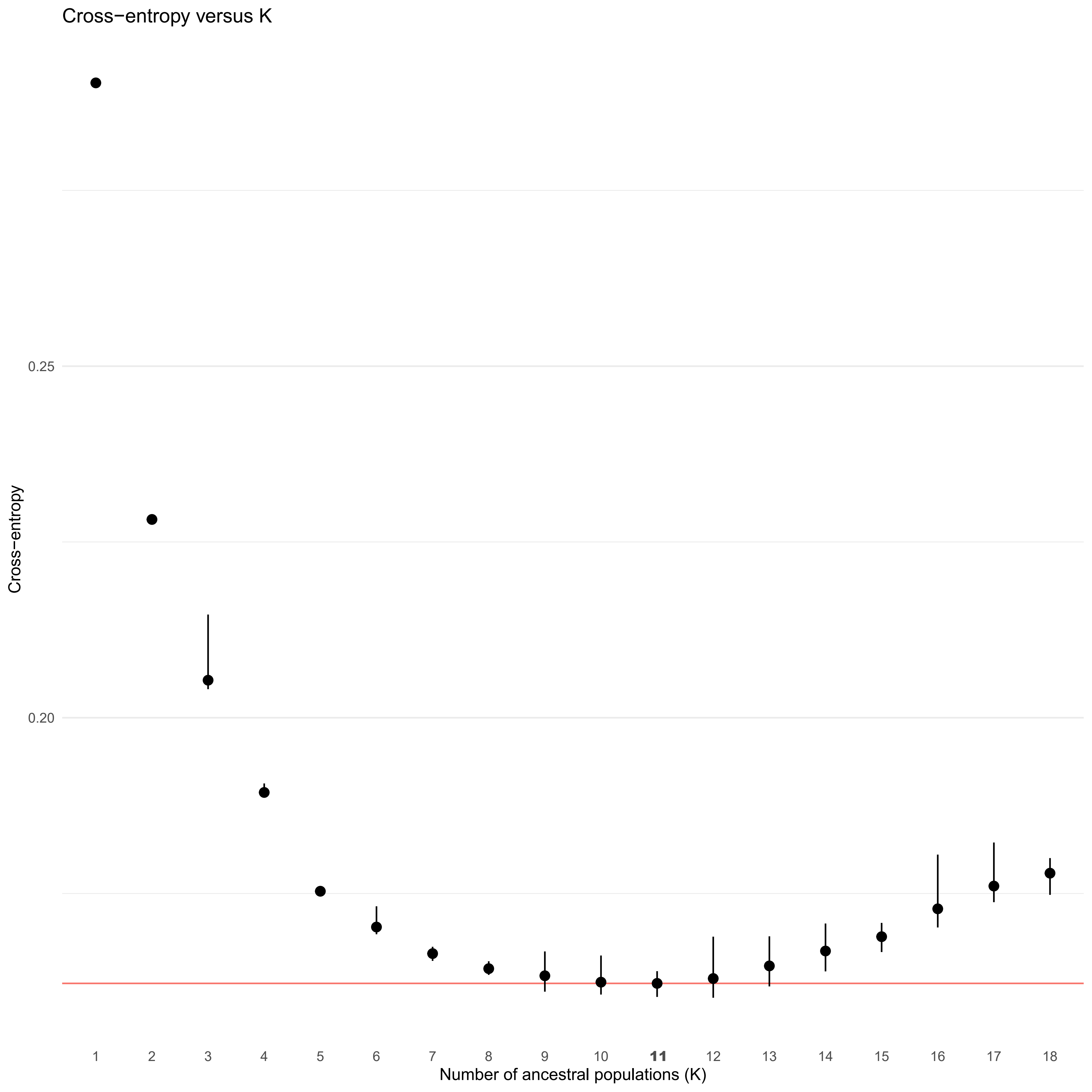

### Fig. S4

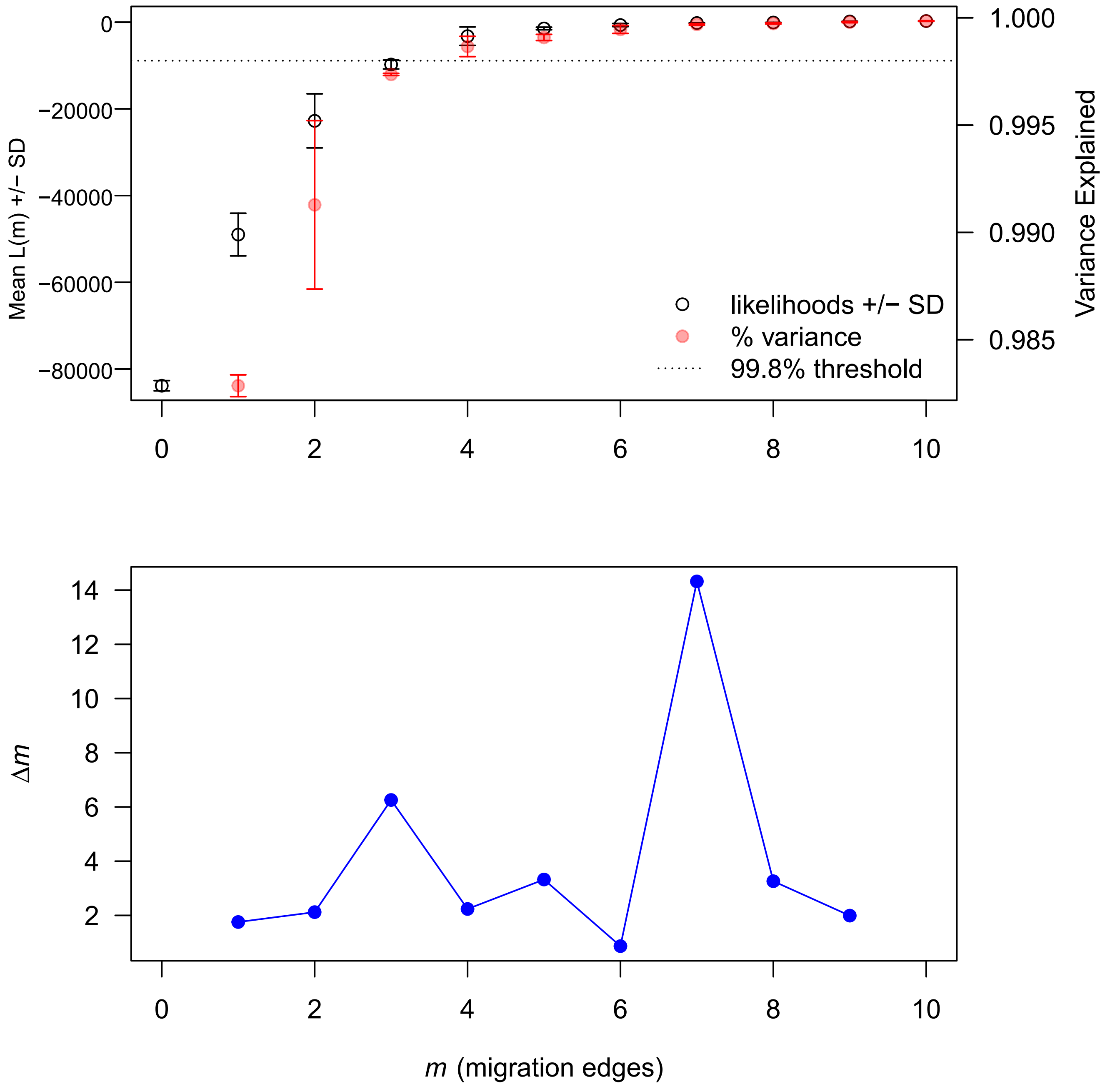

### Fig. S5

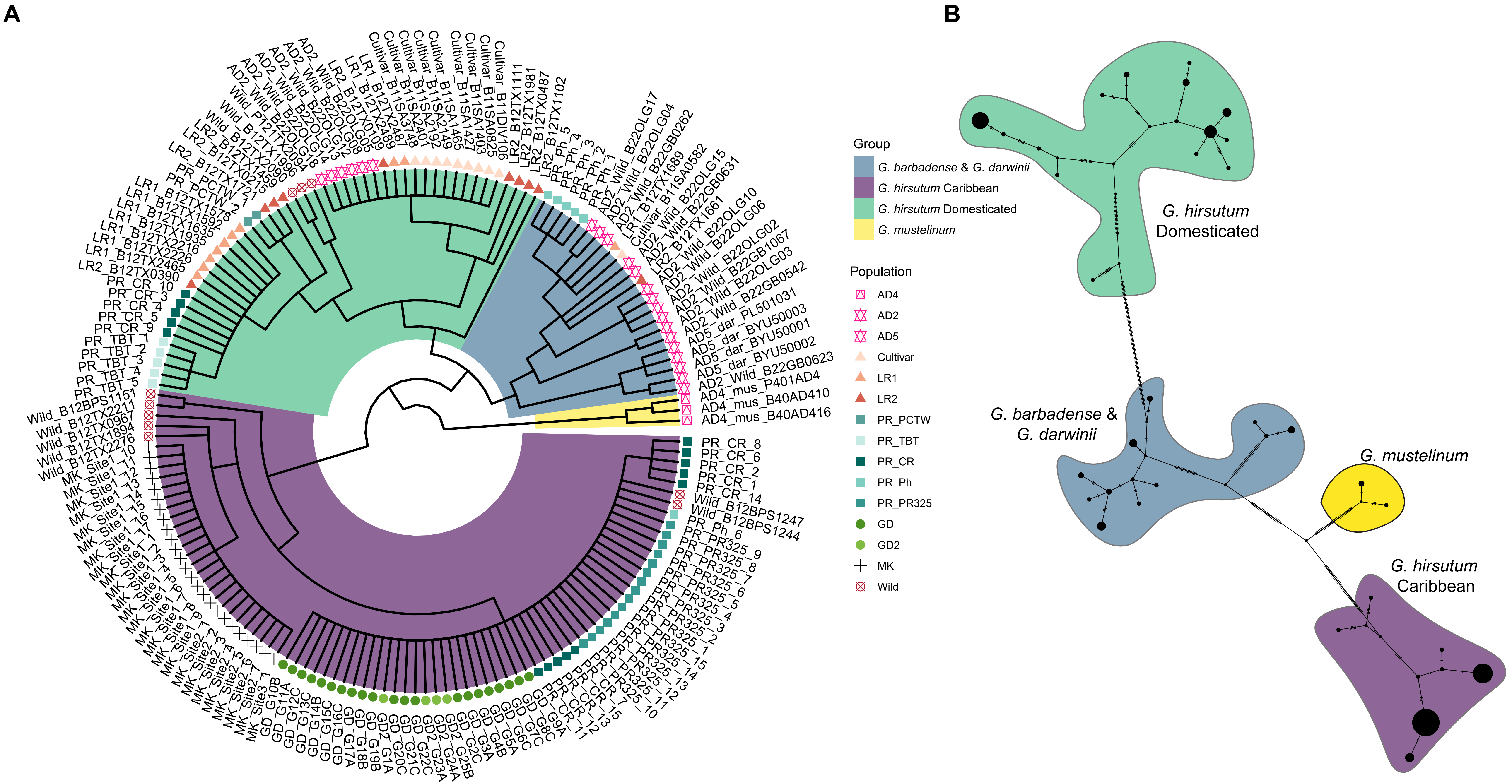

### Fig. S6

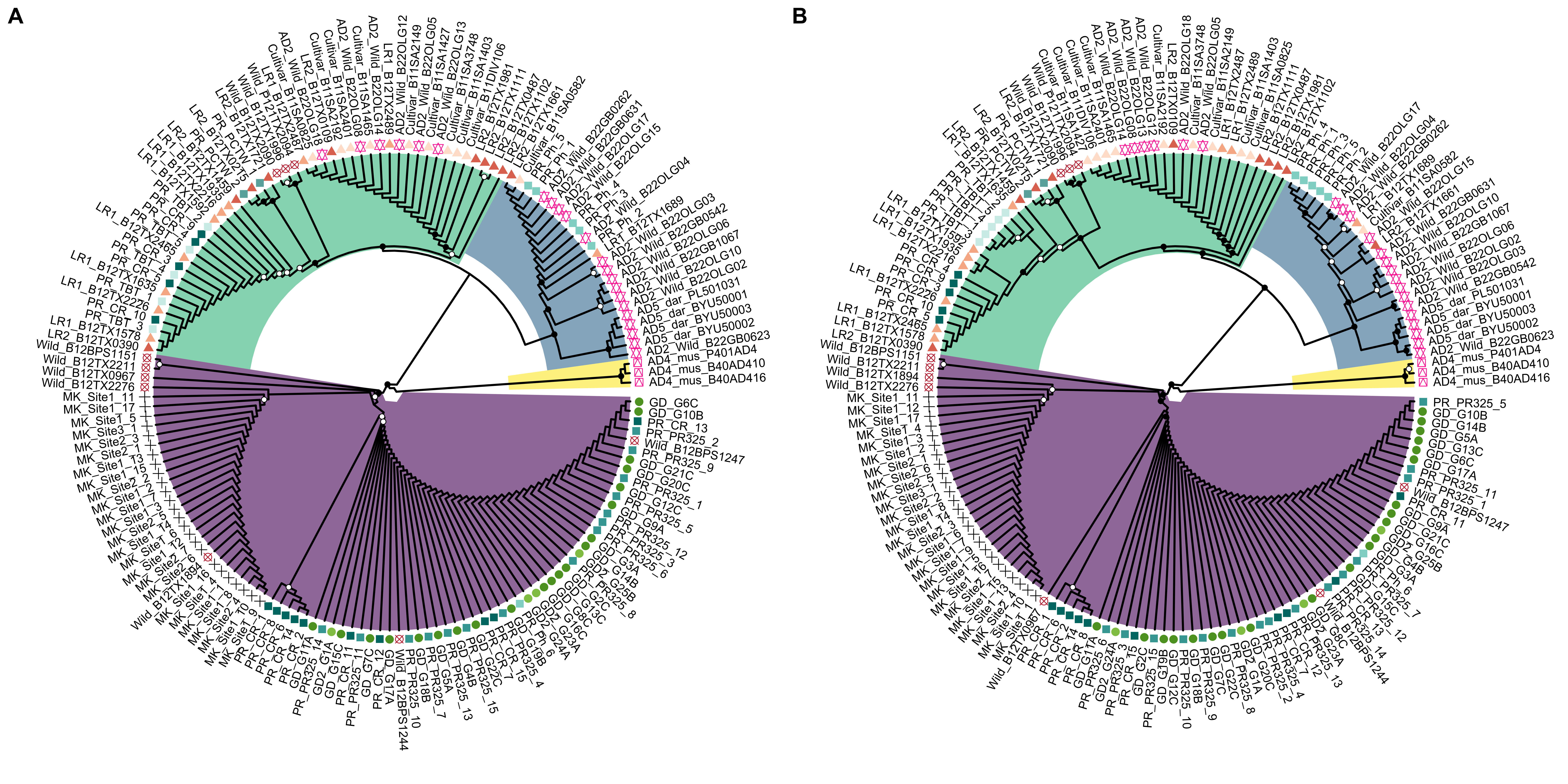

### Fig. S7

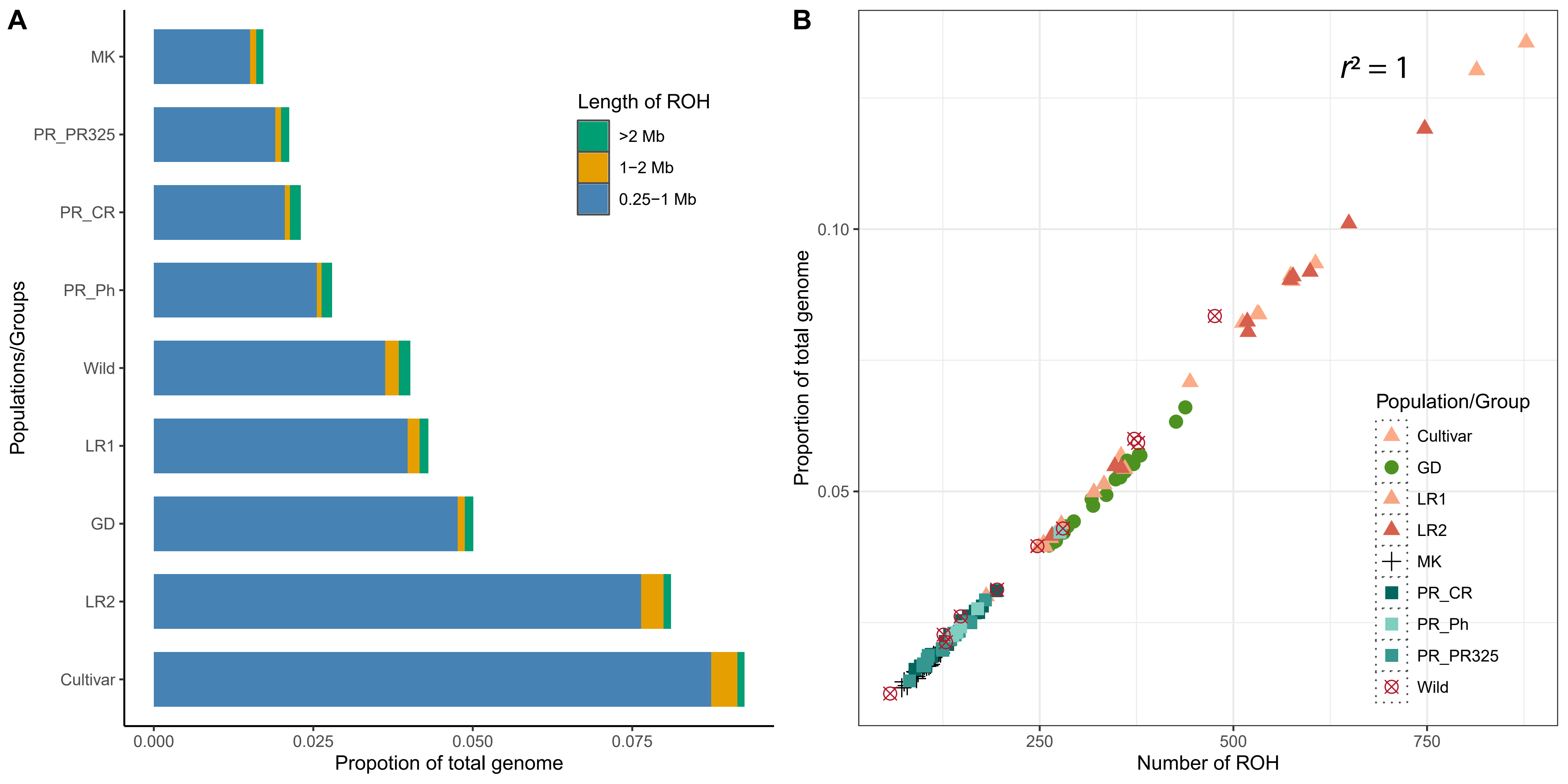

### Fig. S8

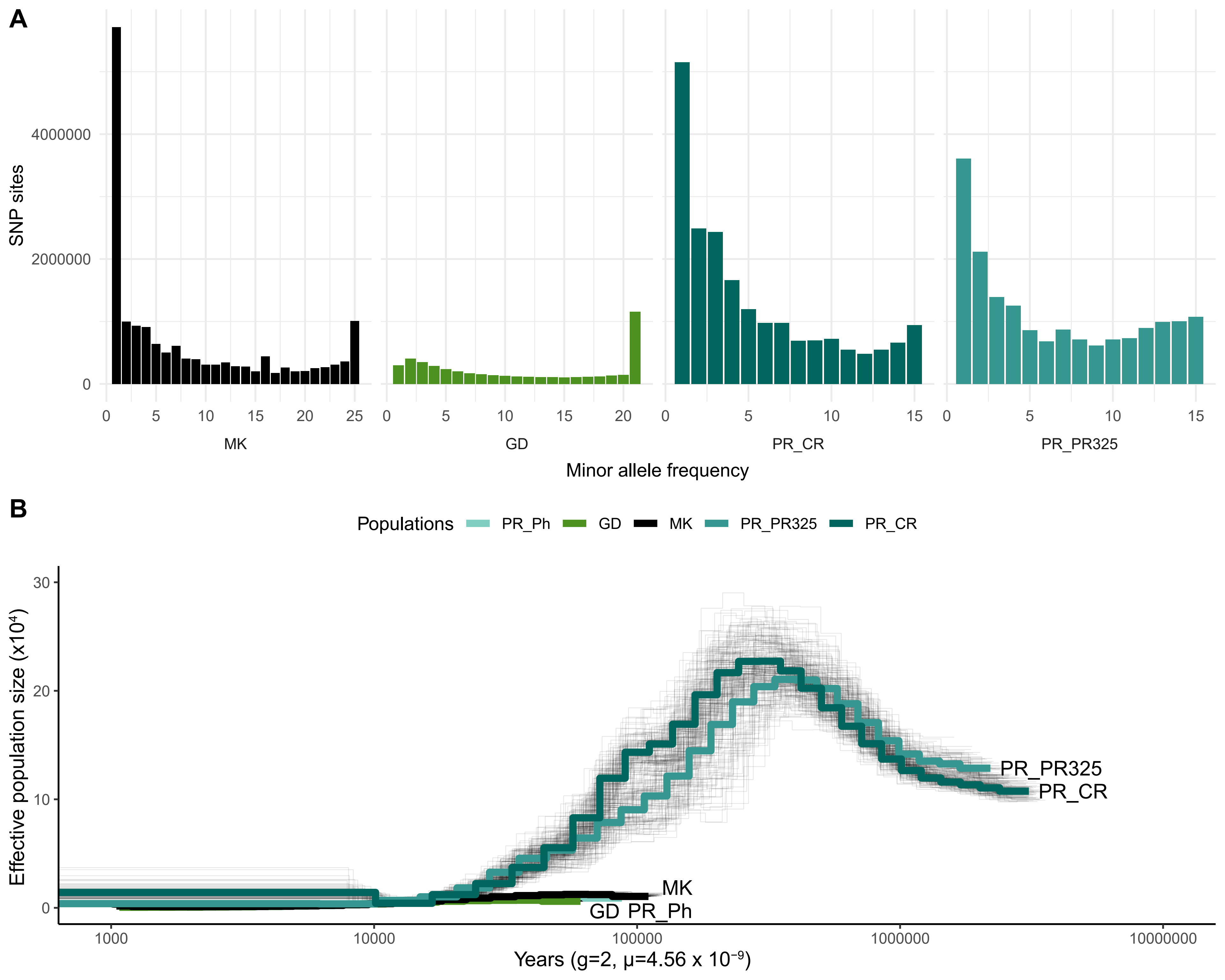
